# Environment-responsive individual cell growth behavior shapes stochastic and deterministic population establishment in ammonia-oxidizing bacteria

**DOI:** 10.64898/2026.05.07.723170

**Authors:** Shuto Ikeda, Hirotsugu Fujitani, Satoshi Tsuneda

## Abstract

Many environmental bacteria do not readily grow under laboratory conditions and population establishment often occurs stochastically. Although the scout hypothesis has been proposed to explain stochastic population establishment in environmental bacteria, how stochastic population establishment is shaped by individual cell growth behaviors in environmental isolates remains unclear. In the present study, we focused on the ammonia-oxidizing bacterium *Nitrosomonas* sp. PY1 and showed that environmentally responsive individual cell growth behavior, incorporating time-dependent stochastic growth initiation, shapes both deterministic and stochastic population establishment dynamics. Using single-cell observation, we revealed that PY1 altered cell growth behavior in response to surrounding biomass production (Δ*V_t_*). These Δ*V_t_*-dependent changes in growth behavior were suppressed by the addition of its own cell-free supernatant (CFS), indicating the presence of a growth regulation mechanism via cell–cell communication. Replicate cultures under the same conditions showed that the population establishment of PY1 was stochastic, whereas the model strain *Nitrosomonas europaea* exhibited synchronized population establishment, consistent with previous reports. This stochasticity in PY1 was also eliminated by the addition of CFS. Finally, a simulation model based on Δ*V_t_*-dependent cell growth behavior of PY1 successfully reproduced synchronized population establishment in the presence of CFS. By contrast, the stochastic population establishment observed in the absence of CFS was successfully reproduced by a model incorporating Δ*V_t_*-independent growth initiation following a Weibull distribution. Such environmentally responsive changes in population establishment dynamics may contribute to the low isolation success of environmental bacteria and sudden blooms of the rare biosphere.

## Introduction

Many environmental bacteria do not readily form colonies on standard agar plates [1]. This phenomenon has been attributed to non-proliferative states such as the viable but non-culturable (VBNC) state and dormancy [1–3]. These states enable bacteria to survive under unfavorable conditions, thereby forming a seed bank that contributes to microbial diversity [4–6]. Moreover, the rare biosphere could influence community structure and stability, implying that bacteria remaining non-proliferative or at low abundance may play a key role in shaping microbial communities [7, 8]. Therefore, elucidating the growth dynamics of non-proliferative environmental bacteria is important for understanding microbial community dynamics.

In some environmental bacteria, resuscitation- and growth-promoting substances, such as resuscitation-promoting factor (rpf) and peptide fragments, can stimulate resuscitation and growth of bacteria from non-proliferative states [9–17]. Based on these findings, the scout hypothesis has been proposed as a growth strategy for environmental bacteria, in which scout cells stochastically initiate growth and subsequently induce the growth of surrounding cells [2, 18, 19]. Supporting this hypothesis, stochastic population establishment has been reported in environmental samples cultured under laboratory conditions [20, 21]. Furthermore, DNA barcoding analyses and single-cell observations of model bacteria have shown that resuscitation from dormancy can occur stochastically [22–24]. However, individual cell growth behavior of environmental isolates has rarely been investigated, and how stochastic population establishment emerges from individual cell growth behavior in environmental bacteria remains largely unexplored.

Ammonia-oxidizing bacteria (AOB) are a group of environmental bacteria known to exhibit stochastic growth behavior. AOB play a crucial role in the global nitrogen cycle; however, their low cultivation success after dilution to extinction has long been recognized as a major challenge in their isolation and cultivation [25, 26]. Furthermore, in some strains of *Nitrosomonas* that have been successfully isolated, nitrite production becomes heterogeneous after subculturing, even though culture conditions such as temperature, pH, and medium composition are optimized [27]. As nitrite production serves as a population-level growth indicator for AOB, some *Nitrosomonas* strains have been interpreted as exhibiting population-level growth heterogeneity [28, 29]. Empirically, subculturing with 10% of late-exponential phase culture supernatant has been reported to promote the stable maintenance of AOB cultures [30]. However, individual cell growth behavior of AOB in the presence and absence of culture supernatant has not yet been quantitatively characterized. Consequently, the cellular basis of the population-level growth heterogeneity in AOB remains unclear.

To address these gaps, we established single-cell observation and quantitative image analysis methods for *Nitrosomonas* and quantified individual cell growth behaviors, including generation time and elongation rate. We further quantified the population establishment dynamics under conditions that resulted in either stochastic or deterministic outcomes. Finally, we constructed a cell elongation and division model based on individual cell growth behaviors, which successfully reproduced the population establishment dynamics. By integrating single-cell observations with modeling, this study links individual cell growth behavior to population establishment in *Nitrosomonas* and provides a quantitative framework for understanding stochastic population establishment.

## Methods

### Strains and culture conditions

*Nitrosomonas* sp. PY1, isolated from activated sludge, and *Nitrosomonas* europaea NBRC 14298, obtained from the Biological Resource Center (NBRC), National Institute of Technology and Evaluation (NITE), were used [26].

PY1 was cultured in batch under dark conditions at 23°C using an inorganic medium containing 2 mM NH_4_Cl, 2.5 mM NaHCO_3_, and 500 U mL^-1^ catalase (FUJIFILM Wako, Japan) [30, 31]. *N. europaea* was cultured in batch under dark conditions at 28°C using 829 medium (HEPES-buffered) according to the protocol provided by NBRC. Detailed culture procedures are described in the Supplementary Methods.

Cell-free supernatant (CFS) was prepared by sequential filtration of exponential-phase cultures of PY1 through a 0.2 µm and 0.22 µm filter (PTFE and MCE, respectively; Millipore, USA), based on a previously described method [10, 17]. CFS was freshly prepared from the preculture samples used in each experiment. The preculture samples contained up to 2 mM ammonia and nitrite; however, as CFS was added at a ratio of 10% (v/v), these components were sufficiently diluted and were unlikely to affect the experimental results.

### Time-lapse imaging

A microfluidic device was constructed according to a previously described procedure [32]. The PDMS microfluidic chips were provided by Professor McKinney. Processing of cover glasses (Thermo Fisher Scientific, USA) and preparation of cellulose membranes (Repligen, USA) were performed as described previously [32, 33] (see the Supplementary Methods for details). Cellulose membranes with a molecular weight cut-off of 12–14 kDa were used.

Cultures of PY1 and *N. europaea* were concentrated 100-fold and 20-fold, respectively, using a 0.2 µm PTFE filter, and cells were subsequently harvested by centrifugation (18,870 × g, 10 min, 23°C). The harvested cells were resuspended in fresh medium to OD_600_ = 1.75. A 5 µL aliquot of the cell suspension was spotted onto the cellulose membrane, and the device was assembled.

Prior to the start of observation, the medium was perfused at a flow rate of 32 µL min^-1^ for approximately 24 h using a syringe pump SPE-1 (AS ONE, Japan) to remove excess cells. At the start of observation, the medium perfusion rate was changed to 8 µL min^-1^. Immediately after assembly through the observation period, the device was kept in the dark at 23°C (PY1) or 28°C (*N. europaea*). The strain-specific culture medium described above was used. When CFS was added, it was supplemented at 10% (v/v) relative to the original medium, based on a previous report [30]. Observations were performed in biological replicates (n = 3).

Time-lapse imaging was performed using an inverted microscope with a 100× oil immersion objective. Detailed microscope configurations are provided in the Supplementary Methods.

### Image processing and cell tracking

Image processing followed a sequential pipeline. First, image correction was performed using Fiji [34], followed by cell segmentation using the StarDist model and the StarDist-OPP model [35–37]. Subsequently, cell tracking was conducted using custom Python code. Finally, cell lineage construction was performed using TrackMate [38]. The StarDist model was trained using manually annotated images based on previously described procedures [36, 39]. All cell lineages were manually inspected and corrected. Detailed image-processing procedures, tracking criteria, and exclusion rules are described in the Supplementary Methods.

### Extraction of individual cell growth parameters

Following a previous study [40], the eight parameters were calculated for each tracked cell: time at birth (*Tb*), time at division (*Td*), generation time (𝑇), area at birth (𝐴𝑏), area at division (𝐴𝑑), elongation rate (𝛼), cell area elongation until division (Δ𝐴), and division ratio (𝐷𝑅) (Fig. S1).

*Tb* was defined as the frame in which cell division was detected. Cell division was identified using an adjustable watershed algorithm (tolerance = 3.0) [41]. If the daughter cell could not be detected immediately after division, the nearest frame detected after division was defined as *Tb*. *Td* was defined as the frame immediately preceding division. If the cell could not be detected immediately before division, the nearest frame detected before division was defined as *Td*. The generation time, 𝑇, was defined as the time from cell birth to division, as 𝑇 = 𝑇𝑑 − *Tb*.

𝐴𝑏, 𝐴𝑑, and 𝛼 were estimated by least-squares fitting of each cell’s area trajectory using the following exponential growth model (Fig. S2):

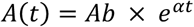

where 𝐴(𝑡) represents the cell area at time 𝑡. To focus the analysis on exponentially growing cells, cells with a coefficient of determination (R^2^) ≤0.75 were excluded from the analysis. 𝐴𝑑 was obtained by substituting the generation time 𝑇 into the exponential growth model. Δ𝐴 was defined as Δ𝐴 = 𝐴𝑑 − 𝐴𝑏. 𝐷𝑅 was defined as the ratio of the area of each daughter cell to the combined area of both daughter cells.

### Estimation of biomass production Δ*V_t_*

To quantify biomass production per time interval in the observed field of view (FOV), we defined Δ*V_t_* as the biomass production during the time interval from 𝑡 to 𝑡 + Δ𝑡:

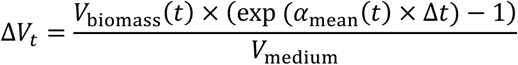

where 𝑉_biomass_(𝑡) is the total cell volume within the FOV at time 𝑡, 𝛼_mean_(𝑡) is the mean elongation rate of cells within the FOV at time 𝑡, Δ𝑡 is the time interval used for the analysis (PY1: 100 min; *N. europaea*: 50 min), and 𝑉_medium_ is the volume of medium flowing into the FOV during Δ𝑡.

Note that Δ*V_t_* is not a directly measured increment of biomass, but rather an estimated value calculated from the instantaneous biomass in the observed FOV and the elongation behavior of the cells. Detailed procedures for calculating the parameters used to derive Δ*V_t_* are described in the Supplementary Methods.

### Batch culture experiments

For batch culture experiments, replicate cultures were prepared in well plates following previously described procedures [27]. Cells were inoculated into a 48-well plate (CORNING, USA) at initial cell densities of 10^5^, 10^3^, and 10^1^ cells mL^-1^. Each well contained 1 mL of medium and 12 replicates were prepared for each condition. To prevent evaporation, the outer wells were filled with sterile medium and were not used in the experiments. Cultures were incubated in the dark at 23°C (PY1) or 28°C (*N. europaea*). The strain-specific media described above were used. When CFS was added, it was supplemented at 10% (v/v), based on a previous report [30]. Experiments were performed in biological replicates (n = 3).

For cell counting, an aliquot of culture was spotted onto a 12-well glass slide (Matsunami Glass Ind., Japan) and air-dried for at least 30 min, followed by staining with SYTOX Green (Thermo Fisher Scientific, USA). Five FOVs per well were imaged, and cell density was determined based on three wells.

NO_2-_ was quantified by the Griess assay using a fluorescence plate reader Synergy H1 (Agilent, USA) until it reached 1.0 mM [42]. The time at which NO_2_^-^ reached 0.25 mM was defined as the timing of population establishment based on linear interpolation.

### Simulation model

An individual-based model (IbM) was constructed in which each cell independently underwent elongation and division. To simplify the model, spatial structure and cell–cell interactions were not considered. The model parameters were obtained through parameter estimation and regression modeling based on single-cell observations of PY1 in the absence of CFS (Fig. S9).

As shown in Fig. 5A, cells were assumed to elongate at each time step and divide upon satisfying the defined conditions. Initial cells were assigned the following parameters: initial cell area (𝐴_0_), initial elongation rate (𝛼_0_), initial generation time (𝑇_0_), minimum cell area required for division (𝐴_min,div_), and division ratio (𝐷𝑅). Cells elongated exponentially and divided according to 𝐷𝑅 once the cell area exceeded 𝐴_min,div_ and the generation time exceeded 𝑇_0_. Daughter cells acquired elongation rate (𝛼) and generation time (𝑇) based on the biomass production (Δ*V_t_*) at birth and the area at birth (𝐴𝑏), while 𝐴_min,div_ and 𝐷𝑅 were sampled from normal distributions. Error terms for 𝐴_min,div_, 𝛼, and 𝑇 were modeled as normal distributions, with variance defined as a function of Δ*V_t_* based on residuals from the regression model.

To reproduce individual cell growth behavior within the microfluidic device, a continuous culture system was assumed and metabolite accumulation was neglected. To simulate population establishment in batch cultures, metabolite accumulation was incorporated, including Δ*V_t_* and nitrite. To represent the decrease in growth rate of PY1 associated with nitrite accumulation, an apparent inhibition constant (𝐾*_i_*) of 100 µM was applied, and the cell elongation rate was limited according to a non-competitive inhibition model. This 𝐾*_i_* was not intended to represent inhibition by nitrite itself, but rather a phenomenological parameter encompassing unmodeled factors (Fig. S11). The effect of CFS was incorporated as the initial Δ*V_t_* at the start of culture, calculated from the amount of NO_2^-^_ in the CFS based on the growth yield of PY1 (33.5 cells pmol⁻¹) and the steady-state cell birth volume (approximately 0.48 µm^3^)[31].

Simulations were performed using Python 3.11.14, and figures were generated using Python 3.11.14 and R 4.4.1. Details of the model, parameter settings, and evaluation methods are described in the Supplementary Methods. The simulation code is available on GitHub (https://github.com/shmy-04/aob-stochastic-growth). The simulation code was developed with the assistance of ChatGPT-5, and the model design, parameter settings, and interpretation of the results were performed by the authors.

## Results

### Distinct individual cell growth behavior between *Nitrosomonas* sp. PY1 and *N. europaea*

*Nitrosomonas* sp. PY1 and *N. europaea* were selected as the strains showing heterogeneous and uniform population establishment timing, respectively. To analyze individual cell growth behavior, a microfluidic device was constructed following previously described procedures [32]. The device enables continuous medium supply while preventing cell washout by sandwiching cells between a cover glass and a cellulose membrane.

Time-lapse imaging revealed dynamic changes in individual cell growth behavior in PY1 (Movie S1). Each cell was segmented using a custom-trained StarDist model [35] (Fig. 1A; see the Methods). The area trajectories were fitted to an exponential growth curve, and individual cell growth curves were obtained for cells with R² > 0.75 (Fig. 1B, Fig. S2). For PY1, cells born during the early observation period (< 60 h) showed prolonged generation times and larger cell areas at division (Fig. 1C). By contrast, cells born during the late observation period (> 60 h) exhibited shorter generation times and divided at smaller cell areas. These temporal changes in generation time and cell area were not observed in *N. europaea* (Fig. 1D).

**Figure 1.**
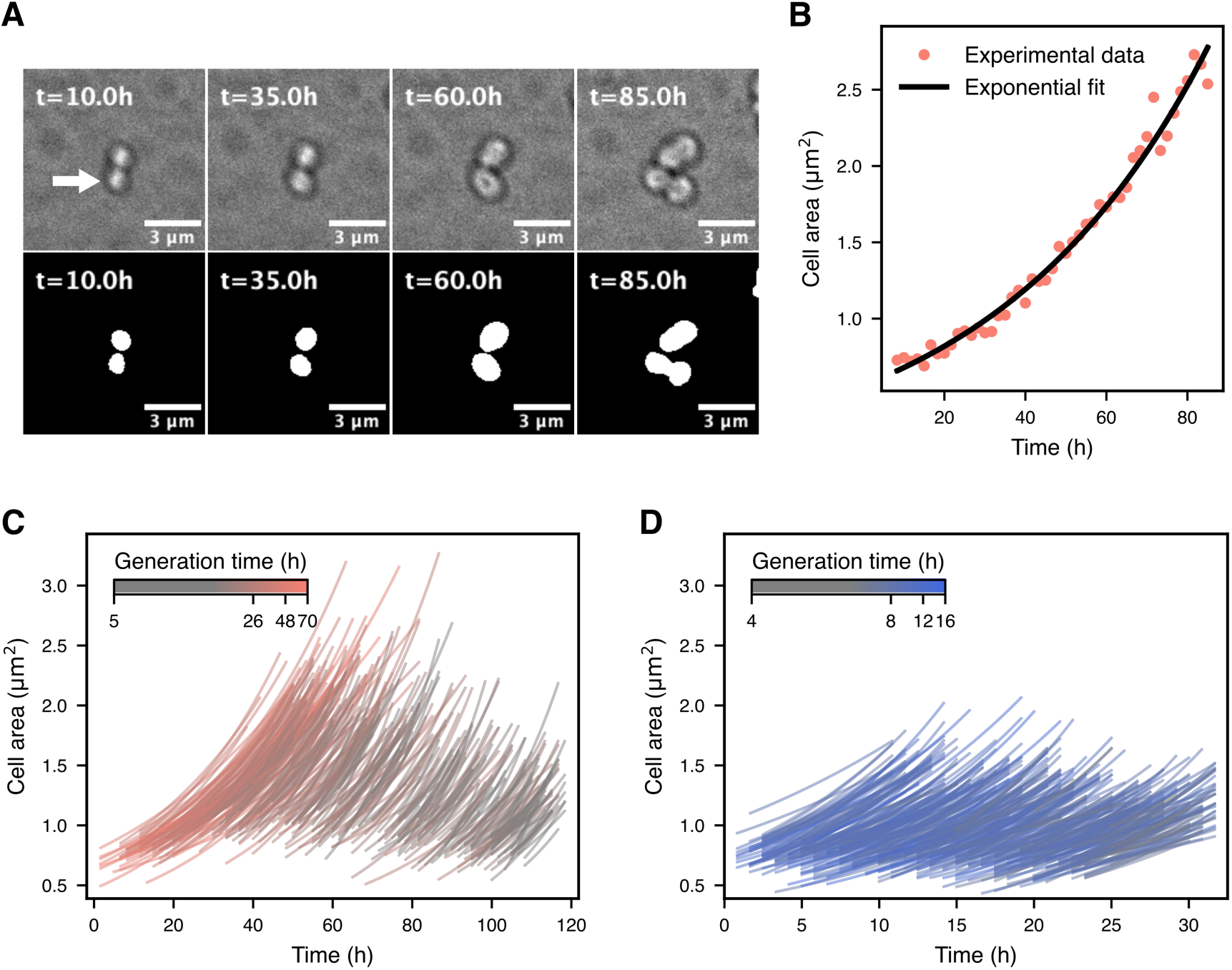
Individual cell growth behavior of two *Nitrosomonas* strains. (A) Representative time-lapse images of a PY1 cell (t = 10.0–85.0 h). The upper panels show images of the cell indicated by the white arrow, from immediately after birth (10.0 h) to immediately before division (85.0 h). The lower panels show the corresponding segmentation masks of the same field of view (FOV) generated using a custom-trained StarDist model [35]. (B) Exponential fit of the cell area trajectory from birth to division for the cell indicated in (A). The same fitting was applied to all tracked cells, and only cells with R² > 0.75 were used in subsequent analyses (PY1: 3747/5325 cells; *N. europaea*: 3850/5320 cells). (C, D) Individual cell growth curves from a representative FOV for PY1 (C) and *N. europaea* (D). Cell area was calculated by multiplying the number of pixels for each cell by the area per pixel (0.0645 × 0.0645 µm^2^). For visualization, the observation period was divided into 20 bins, and 20 cells were randomly sampled from each bin. Line color indicates the generation time of each cell.

### Δ𝑉𝑡-dependent modulation of individual cell growth behavior in *Nitrosomonas* sp. PY1

Because cells were not washed out from the microfluidic device, the biomass within the observed FOVs increased over time (Fig. S3A, D). In PY1, the mean elongation rate of cells within the FOVs also increased over time (Fig. S3B), indicating a rapid increase in instantaneous biomass production. By contrast, *N. europaea* exhibited a high elongation rate from the early observation period, and the mean elongation rate either remained constant or decreased over time (Fig. S3E). Based on these results, we hypothesized that while *N. europaea* competes with surrounding cells for available substrates, PY1 may increase its own growth activity as the surrounding cells become more metabolically active. To test this hypothesis, we calculated the biomass production in the time interval from 𝑡 to 𝑡 + Δ𝑡 (Δ*V_t_*) for each observed FOV (Fig. S3C, F; see the Methods).

We analyzed the relationship between Δ*V_t_* at cell birth and individual cell growth parameters: generation time (𝑇), elongation rate (𝛼), and area elongation until division (Δ𝐴) (Fig. 2, Fig. S1). In PY1, 𝑇 of cells born under low Δ*V_t_* conditions were substantially prolonged (r = -0.52, Fig. 2A). Similarly, 𝛼 varied with Δ*V_t_*; however, the change in 𝛼 was less pronounced than that of 𝑇 (r = 0.42, Fig. 2B). Consistent with 𝑇, Δ𝐴 decreased substantially with Δ*V_t_* (r = -0.57, Fig. 2C). In *N. europaea*, no clear Δ*V_t_*-dependent modulations in growth behavior, such as those observed in PY1, were detected (Fig. 2D–F). These results indicate that PY1 exhibits Δ*V_t_*-dependent changes in individual cell growth behavior, whereas *N. europaea* maintains stable growth behavior.

**Figure 2.**
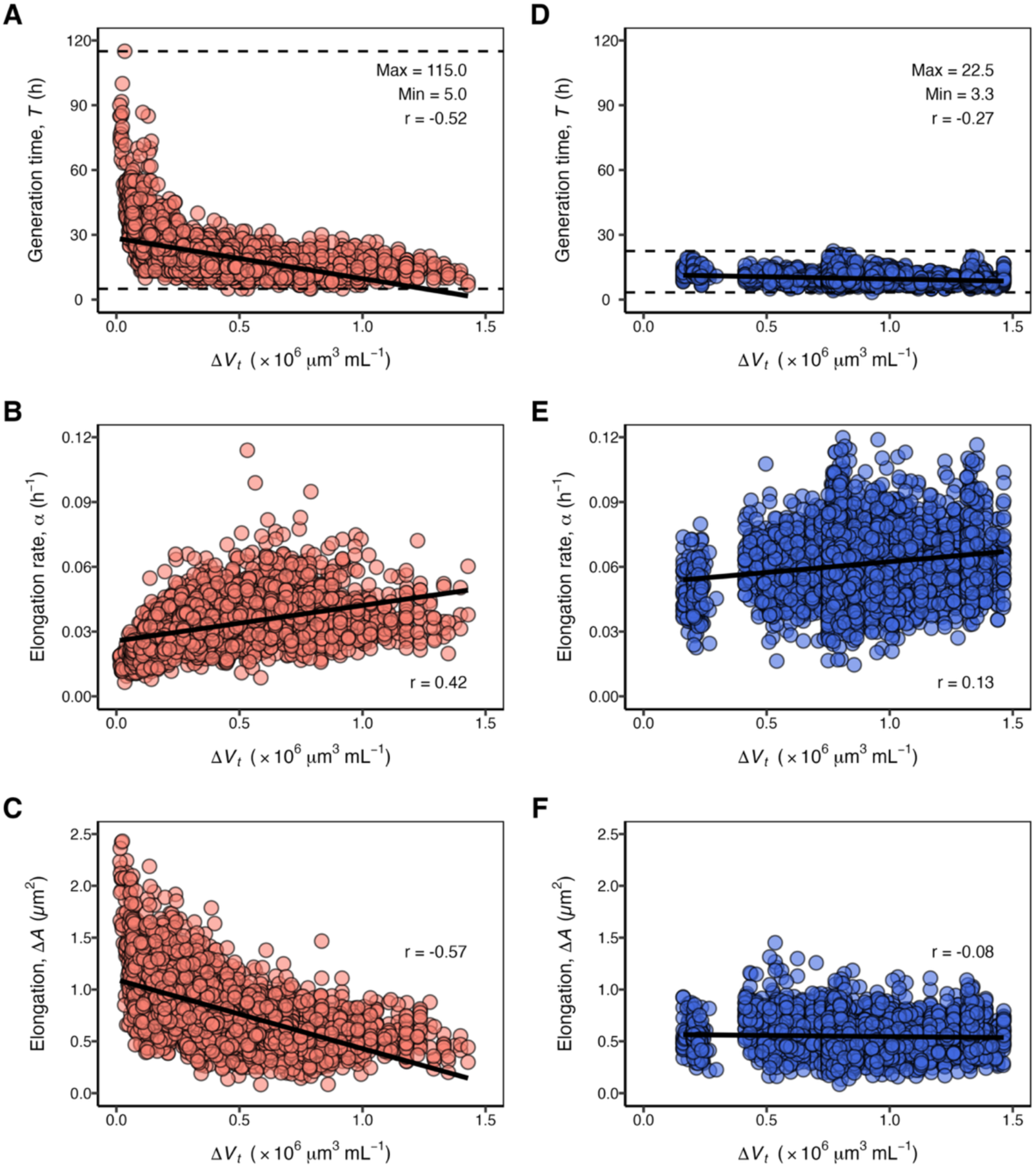
Relationships between individual cell growth parameters and biomass production (Δ𝑽_𝒕_) in two *Nitrosomonas* strains. Individual cell growth parameters for PY1 (A-C) and *N. europaea* (D–F). (A, D) Generation time (𝑇) as a function of Δ*V_t_* at cell birth. Dotted lines indicate the maximum and minimum observed generation times. A linear regression line fitted to all data is shown to indicate the overall trend. (B, E) Elongation rate (𝛼) as a function of Δ*V_t_* at cell birth. (C, F) Cell area elongation until division (Δ𝐴) as a function of Δ*V_t_* at cell birth. Three FOVs were analyzed for each time-lapse observation experiment, and data from biological replicates (n = 3) were combined.

### CFS-mediated cell–cell communication modulates growth behavior in *Nitrosomonas* sp. PY1

We investigated which physiological factors are reflected in Δ*V_t_*. Given that many bacteria engage in cell–cell communication mediated by metabolites [43], we performed time-lapse observations of PY1 using CFS derived from the same strain, which is expected to contain cell-derived metabolites (Movie S3).

Compared with the condition without CFS, CFS promoted active cell division from the early observation period and led to smaller cell areas (Movie S4). Visualization of the individual cell growth curves over time showed that changes in generation time and cell area were reduced relative to the condition without CFS (Fig. 3A). Regarding 𝑇, prolongation was still observed under low Δ*V_t_* conditions; however, the maximum generation time decreased from 115 h without CFS to 65 h with CFS, indicating that the Δ*V_t_*-dependent change was substantially reduced (Fig. 3B). By contrast, 𝛼 showed no clear difference compared with the condition without CFS (Fig. 3C). Consequently, the Δ*V_t_*-dependent change in Δ𝐴 was also markedly reduced, similar to the change observed in 𝑇 (Fig. 3D). These results suggest that CFS in the supplied medium partially reproduces the effect reflected by Δ*V_t_* and regulates the growth behavior of PY1.

**Figure 3.**
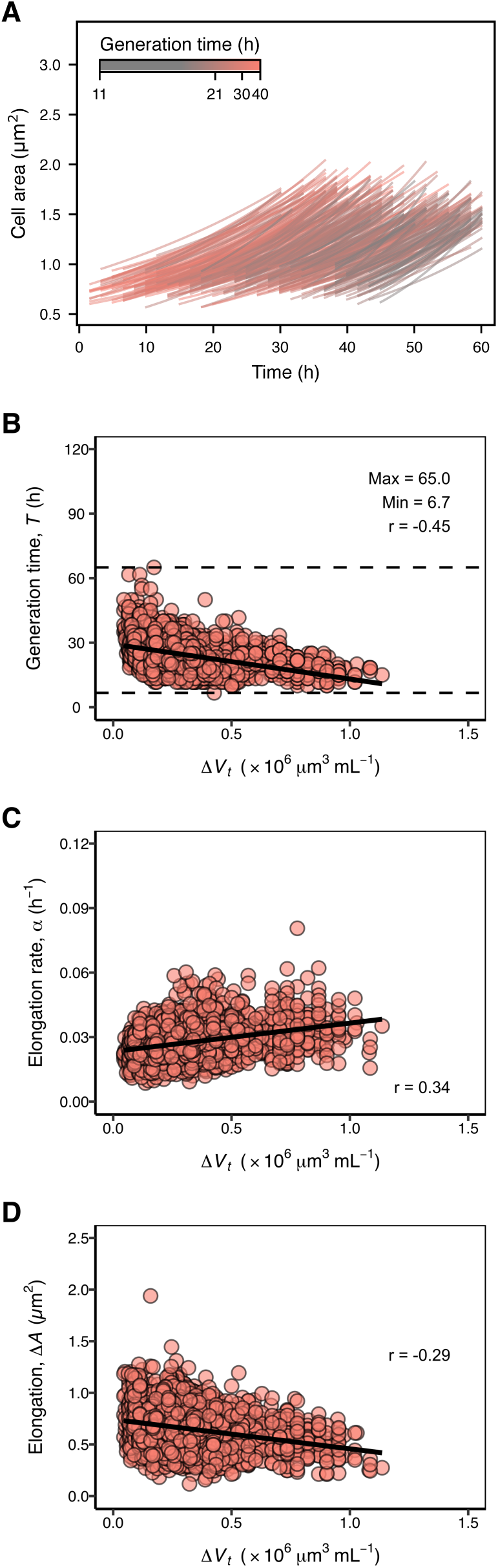
Relationships between individual cell growth parameters and biomass production (Δ𝑽_𝒕_) in PY1 with CFS. (A) Individual cell growth curves from a representative FOV. Individual cell area trajectories from birth to division were fitted to an exponential growth curve. Only cells with R^2^ > 0.75 were used in the analysis (3222/4249 cells). As in Fig. 1C, cell area was calculated from pixel counts, and 20 cells were randomly sampled from each of 20 bins. (B–D) Generation time (𝑇), elongation rate (𝛼), and cell area elongation until division (Δ𝐴) as a function of Δ*V_t_* at cell birth. Three FOVs were analyzed for each time-lapse observation experiment, and data from biological replicates (n = 3) were combined.

### CFS suppresses heterogeneity in population establishment timing in *Nitrosomonas* sp. PY1

When PY1 is inoculated into fresh medium without CFS, the timing of population establishment becomes heterogeneous across cultures [27]. Given that CFS suppressed changes in individual cell growth behavior, we hypothesized that it would also homogenize the timing of population establishment. Therefore, we inoculated PY1 into 48 well plates containing medium supplemented with or without CFS and plotted the time course of the proportion of nitrite-positive wells. Wells in which nitrite concentrations reached ≥ 0.25 mM were defined as nitrite-positive wells and used as an indicator of population establishment.

In the absence of CFS, PY1 gradually increased the proportion of nitrite-positive wells, particularly at low initial cell densities (10^3^ and 10^1^ cells mL^-1^). By contrast, *N. europaea* showed a sharp and synchronous increase in the proportion of nitrite-positive wells, regardless of the initial cell density (Fig. 4A). To quantify variability in the time to reach 0.25 mM nitrite in each well, these times were normalized to the mean under each condition. The SD of the normalized time to reach 0.25 mM nitrite was larger in PY1 than in *N. europaea* (Fig. 4B). At an initial cell density of 10^3^ cells mL^-1^, the SD in PY1 was approximately 7-fold higher than that in *N. europaea* on average (Fig. S5). These results indicate that the timing of population establishment in PY1 becomes heterogeneous under low cell density conditions in the absence of CFS. At an initial cell density of 10^1^ cells mL^-1^ in PY1, several wells did not show nitrite production within the culture period (269 days; Fig. S7); therefore, only the nitrite-positive wells were analyzed.

**Figure 4.**
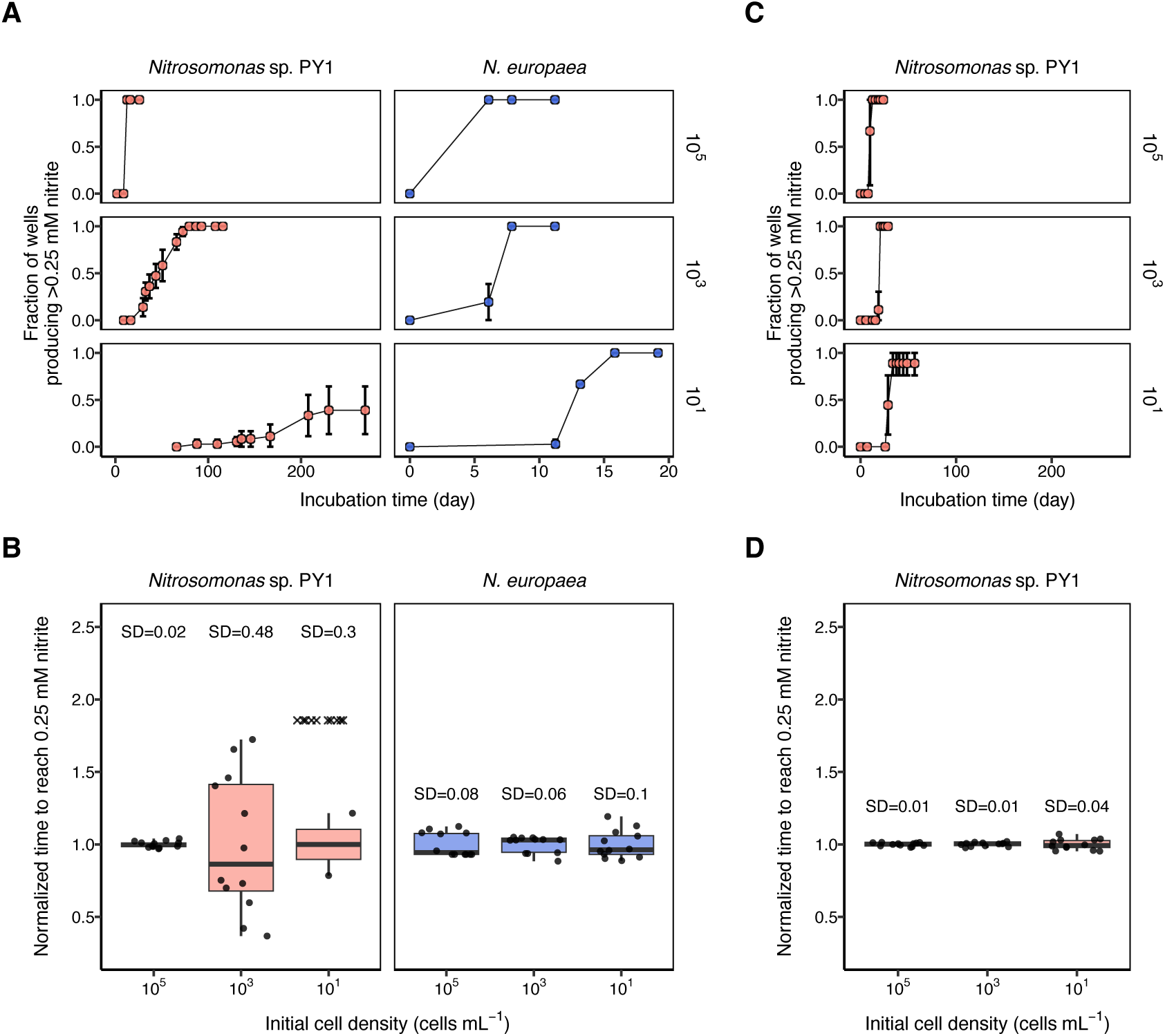
Effects of CFS on heterogeneity in population establishment timing in two *Nitrosomonas* strains. (A) Time course of the proportion of nitrite-positive wells when each strain was inoculated into 48-well plates at 10^5^, 10^3^, and 10^1^ cells mL^-1^ in the absence of CFS. Wells in which nitrite concentration reached ≥0.25 mM were defined as nitrite-positive wells. Experiments were performed in biological replicates (n = 3), and error bars (mean ± SD) are shown. (B) Box plots showing the normalized time to reach ≥0.25 mM nitrite in the absence of CFS. Time to reach ≥0.25 mM nitrite was normalized to the mean under each condition. Wells in which nitrite concentration did not reach the threshold concentration of 0.25 mM during the observation period (269 days) are indicated by crosses (×). Data from a representative replicate are shown. (C) Time course of the proportion of nitrite-positive wells for PY1 at 10^5^, 10^3^, and 10^1^ cells mL^-1^ in the presence of CFS. Experiments were performed in biological replicates (n = 3), and error bars (mean ± SD) are shown. (D) Box plots showing the normalized time to reach ≥0.25 mM nitrite for PY1 in the presence of CFS. Data from a representative replicate are shown.

By contrast, with CFS, PY1 showed a sharp and synchronous increase in the proportion of nitrite-positive wells regardless of initial cell density (Fig. 4C). The SD of the normalized time to reach 0.25 mM nitrite decreased markedly (Fig. 4D). At an initial cell density of 10^3^ cells mL^-1^, the SD in PY1 with CFS was reduced to approximately 1/22 of that without CFS on average (Fig. S6). These results indicate that CFS substantially suppresses heterogeneity in population establishment timing in PY1. Nitrite production was not detected in medium supplemented with CFS alone (Fig. S8).

### IbM reproduces the growth behavior of *Nitrosomonas* sp. PY1 in the microfluidic device

To test whether Δ*V_t_*-dependent individual cell growth behavior could reproduce population-level growth dynamics, an IbM was constructed (Fig. 5A; see the Methods). In this model, each cell was assumed to independently repeat elongation and division. The parameters required for the simulation (𝐴_0_, 𝛼_0_, 𝑇_0_, 𝐴_min,div_, 𝐷𝑅, 𝛼, 𝑇, 𝐴_max_) were obtained from time-lapse observation data of PY1 in the absence of CFS (Fig. S9).

**Figure 5.**
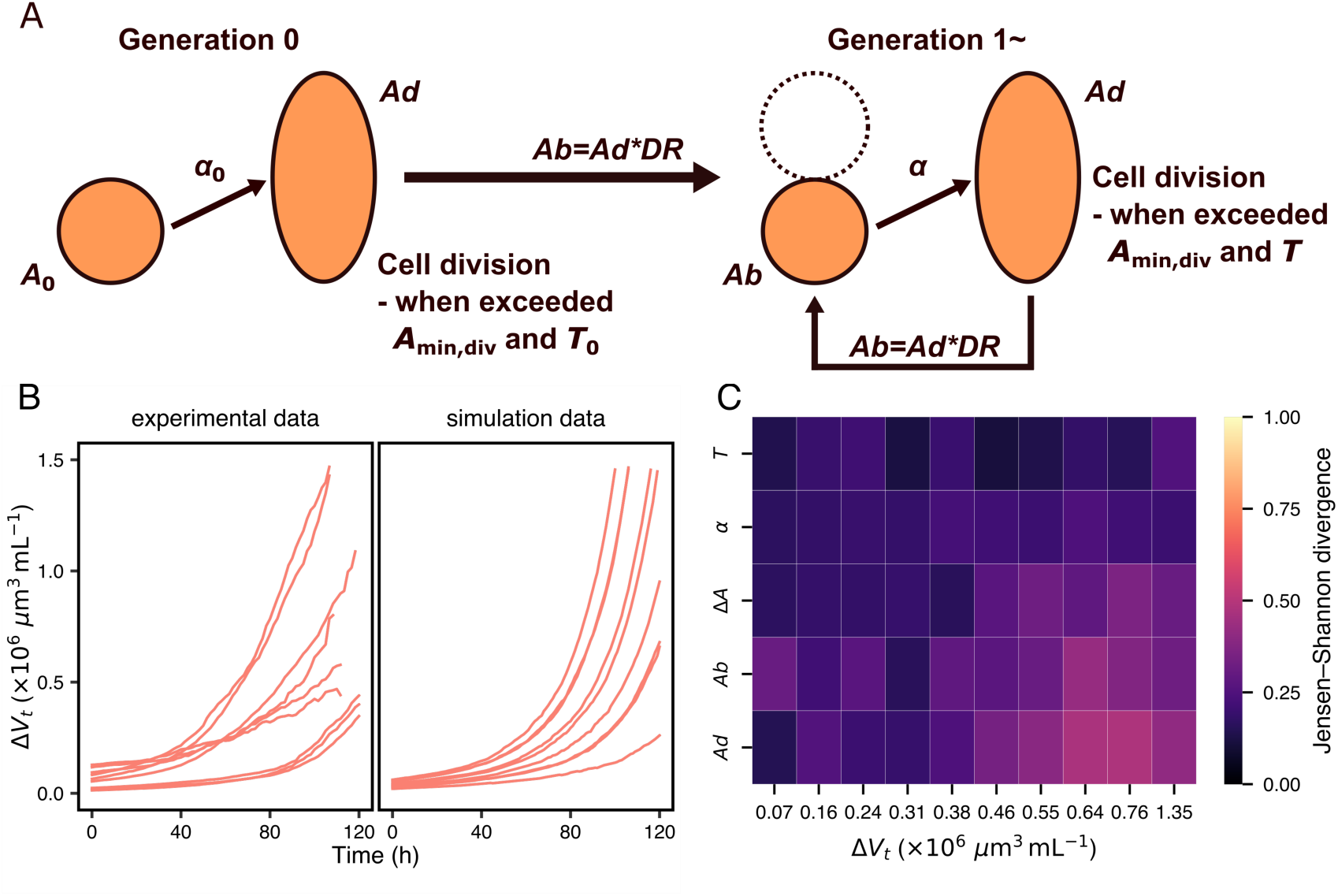
Individual-based modeling of PY1 growth behavior in the microfluidic device. (A) Schematic overview of the model. 𝐴_0_, 𝛼_0_, and 𝑇_0_ represent the initial cell area, elongation rate, and generation time, respectively. 𝐴_min,div_ represents the minimum cell area required for division. 𝐴𝑑 and 𝐴𝑏 represent the cell area at division and at birth, respectively. 𝐷𝑅 represents the division ratio. 𝛼 and 𝑇 represent the elongation rate and generation time in subsequent generations, respectively. (B) Comparison of the time course of Δ*V_t_* between experimental data and simulation. Simulations were performed to match the number of analyzed fields of view (FOVs) (biological replicates, n = 3; 3 FOVs per replicate). The initial number of cells in each simulation was set to match the number of cells successfully tracked in the corresponding FOV (replicate 1: 45, 28, 42; replicate 2: 68, 76, 69; replicate 3: 57, 56, 40 cells). (C) Jensen–Shannon divergence (JSD) of individual cell growth parameters between experimental data and simulation. Individual cell data were binned into 10 groups according to Δ*V_t_* at cell birth, and JSD was calculated for each bin.

The simulations qualitatively reproduced the dynamics of Δ*V_t_* within each FOV (Fig. 5B). Because the simulation assumes only exponentially elongating cells and does not account for non-dividing or growth-arrested cells, we assessed whether the simulation captured overall growth dynamics rather than quantitatively reproducing Δ*V_t_* (Fig. 5B).

To quantify the reproducibility of the distributions of individual cell growth parameters, the experimental and simulation data were partitioned based on Δ*V_t_*, and the Jensen–Shannon divergence (JSD) between the experimental and simulated distributions was calculated for each bin (Fig. 5C). The JSD of all parameters remained ≤ 0.5 in all bins, indicating that the distributions of individual cell growth parameters generated by the simulation were broadly consistent with the experimental distributions. Individual cell growth parameters (𝑇, 𝛼, Δ𝐴, 𝐴𝑏, 𝐴𝑑) as a function of Δ*V_t_* in simulation and experimental data are shown in Supplementary Fig. S10.

### IbM reproduces stochastic population establishment in batch culture

Finally, we tested whether the IbM could reproduce the population establishment dynamics of PY1 in batch cultures (Fig. 4). In the batch culture simulation, accumulation of Δ*V_t_* and nitrite was incorporated, and reproducibility was evaluated based on the time course of the proportion of nitrite-positive wells.

Simulations conducted in the presence of CFS were consistent with the experimental data at initial cell densities of 10^5^ and 10^3^ cells mL^-1^, and 10^1^ cells mL^-1^ (Fig. 6A). At an initial cell density of 10^1^ cells mL^-1^, a slight discrepancy was observed in the timing of appearance of nitrite-positive wells. This discrepancy was reduced using initial cell densities re-estimated from the probability of zero-cell wells in the experimental data, suggesting that the effective initial cell density was lower than the nominal value (Fig. S12).

**Figure 6.**
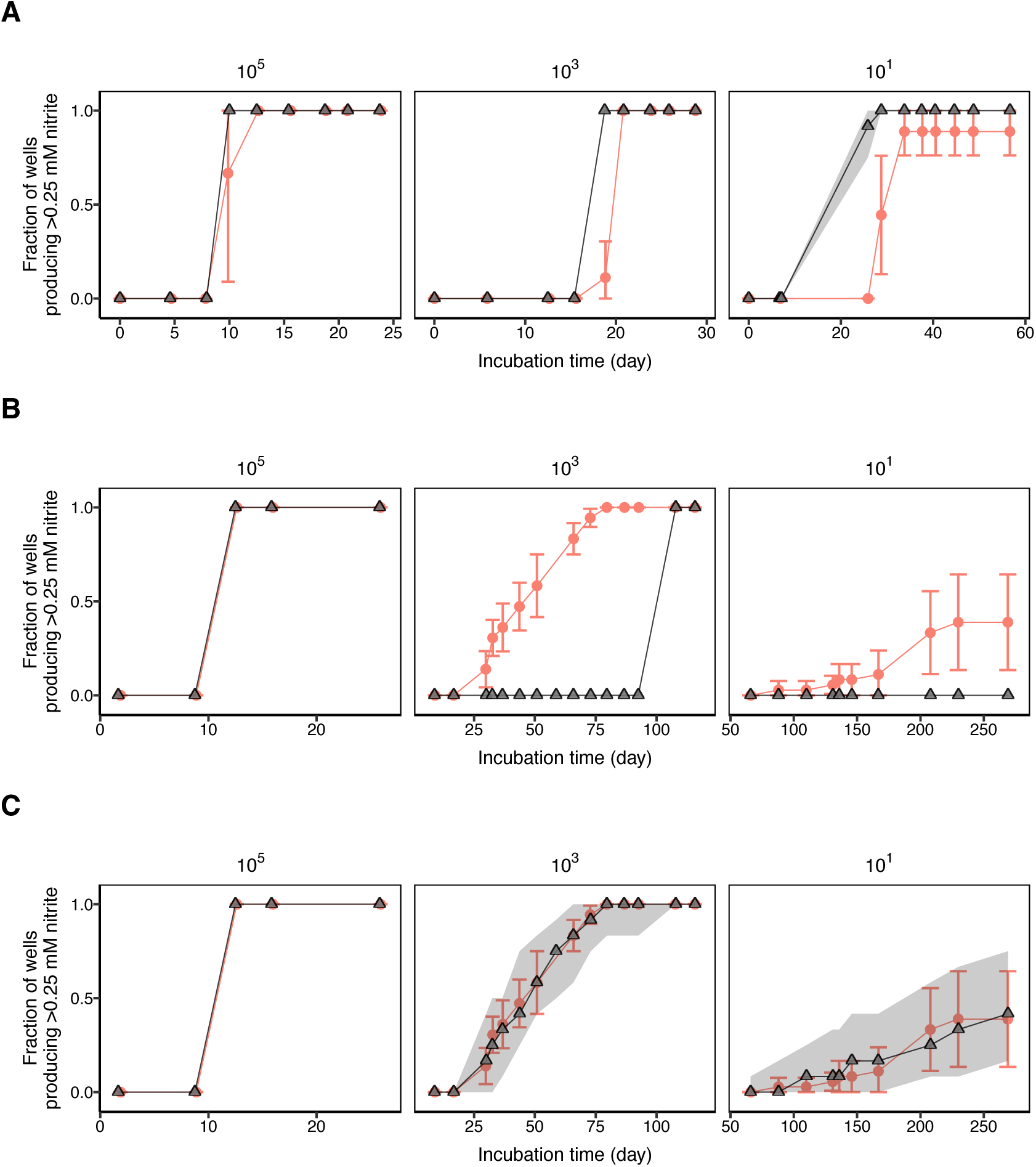
Simulation of nitrite-positive well dynamics for PY1 in batch culture. Growth of PY1 in batch culture was simulated using the model validated in Fig. 5. Nitrite production was calculated from cumulative biomass production (Δ*V_t_*) based on the steady-state cell birth volume and growth yield. Wells in which nitrite concentration reached ≥0.25 mM were defined as nitrite-positive wells. The inhibition constant (𝐾_0_) for nitrite was set to 100 µM. In each simulation, cell growth and nitrite production were simulated for 12 wells, and the proportion of nitrite-positive wells was calculated at each time point. Simulations were performed 100 times independently. The median proportion at each time point is shown, and the 95% prediction interval is indicated by a shaded region. Error bars for experimental data represent mean ± SD. Red circles and lines represent experimental data, and black triangles and lines represent simulation results. (A, B) Experimental data were compared with simulation results in the presence of CFS (A) and in the absence of CFS (B). The initial Δ*V_t_* and nitrite concentrations in the presence of CFS were set according to the corresponding experimental data. (C) Experimental data in the absence of CFS were compared with simulation results incorporating Δ*V_t_*-independent stochastic growth initiation. The probability of growth initiation was modeled using a Weibull hazard function.

In the absence of CFS, the Δ*V_r_*-dependent model alone failed to reproduce the time course of nitrite-positive well proportions at low initial cell densities (10^3^ and 10^1^ cells mL^-1^). When transient biomass production immediately after inoculation was not included, the generation times of initial cells became excessively prolonged. Thus, nitrite-positive wells were not detected under any condition (Fig. S13). Therefore, we assumed transient growth immediately after inoculation of each initial cell, equivalent to 20% of the steady-state birth volume of PY1 cells, based on time-lapse observations (see the Supplementary Methods). Under this assumption, the time course of nitrite-positive well proportions was reproduced at an initial cell density of 10^5^ cells mL^-1^; however, the model did not reproduce the experimental results at initial cell densities of 10^3^ and 10^1^ cells mL^-1^ (Fig. 6B).

These results indicated that Δ*V_t_*-dependent cell growth alone could not explain the population establishment dynamics in the absence of CFS at low initial cell densities. We therefore assumed Δ*V_t_*-independent stochastic growth initiation. For diverse bacteria, including nitrifying bacteria, it has been reported that population establishment probability can be described by the cumulative distribution function (CDF) of growth initiation following a Poisson process with a fixed lag time [44–46]. In this study, the time course of nitrite-positive well proportions fit well with the corresponding CDF only when the parameters were optimized for each initial cell density. However, a single parameter set could not simultaneously fit the experimental data across multiple initial cell density conditions (Fig. S14A, B). In particular, the fit failed to capture the time-dependent increase in the proportion of nitrite-positive wells at an initial cell density of 10^1^ cells mL^-1^. Therefore, we fitted the CDF of a Weibull distribution to the time course of nitrite-positive well proportions to capture time-dependent changes in the probability of Δ*V_t_*-independent growth initiation. Using this approach, time course of the nitrite-positive well proportions at initial cell densities of 10^3^ and 10^1^ cells mL^-1^ fit well with a single set of parameters (shape = 2.12, scale = 1015.2 days; Fig. S14C).

Using the obtained parameters, Δ*V_t_*-independent stochastic growth initiation of individual cells following a Weibull hazard function was incorporated into the model. Consequently, the mean time course of nitrite-positive well proportions in the experimental data fell within the 95% prediction interval of the simulation (Fig. 6C). These results indicate that the population establishment dynamics of PY1 in the absence of CFS can be reproduced by incorporating Δ*V_t_*-independent growth initiation governed by a Weibull distribution.

Note that incorporating Δ*V_t_*-independent growth initiation did not affect the simulation results in the presence of CFS (Fig. S15). The time course of the nitrite concentration in each well in the simulation is shown in Supplementary Fig. S16.

## Discussion

Environmental isolates often exhibit heterogeneity in the timing of population establishment when cultured under laboratory conditions [20, 21, 27], as described by the scout hypothesis [2, 18, 19]. Here, we combined single-cell observations with mathematical modeling to investigate how individual cell growth behavior shapes population establishment dynamics in an environmental isolate, *Nitrosomonas* sp. PY1. PY1 altered its individual cell growth behavior in response to surrounding biomass production (Δ*V_t_*) and CFS, indicating the presence of an environmentally responsive growth regulatory system. By contrast, such Δ*V_t_*-dependent changes were not observed in the model strain *N. europaea* (Figs. 2–4). In the presence of CFS, PY1 exhibited synchronized population establishment, which was reproduced by a Δ*V_t_*-dependent growth model. By contrast, in the absence of CFS, PY1 exhibited heterogeneous population establishment, which was reproduced by a model incorporating Δ*V_t_*-independent stochastic growth initiation (Fig. 6). These findings suggest that, in environmental isolates possessing an environmentally responsive growth regulatory system, active growth in the presence of growth-promoting factors and stochastic growth initiation in their absence can lead to both deterministic and stochastic population establishment. Such growth characteristics may render growth initiation under standard laboratory conditions stochastic, thereby contributing to the low success rate of isolation and cultivation [25, 26, 47–49].

Although catalase supplementation is essential for the cultivation of PY1 [31], population establishment in PY1 remained heterogeneous in the absence of CFS, even when catalase was supplemented (Fig. 4). This suggests that PY1 modulates its growth behavior in response to additional signaling molecules other than catalase. PY1 exhibits a low *Km* value and long generation time compared with other *Nitrosomonas* species [31]. Based on previous findings, PY1 has been considered a K-strategist adapted to oligotrophic environments, characterized by slow growth and efficient substrate utilization under low-nutrient conditions [50]. However, PY1 shortened its generation time in response to CFS to levels comparable to those of the model strain *N. europaea* (Figs. 2 and 3). This suggests that PY1 is not a strict K-strategist, but can adopt an r-strategist-like growth mode, characterized by rapid proliferation, depending on environmental conditions.

Simulations incorporating Δ𝑉_t_-independent stochastic growth initiation following a Weibull distribution reproduced the observed population establishment dynamics in a batch culture in the absence of CFS (Fig. 6C). In a Weibull distribution, the shape parameter determines how the probability of an event changes over time, whereas the scale parameter determines the time scale of the event. The fitted Weibull distribution for PY1 had a shape parameter of 2.12 and a scale parameter of 1015.2 days. Because the shape parameter exceeds 1, the probability of growth initiation in PY1 increases as the lag phase persists. Weibull-distributed lag phase escape has also been reported in *Listeria monocytogenes*, where the shape parameter falls within a range comparable to that of PY1 (1–4) [51, 52]. By contrast, the scale parameter in *L. monocytogenes* was 2–43 h, which was more than 500-fold smaller than that of PY1. These observations suggest that although the time-dependent increase in growth initiation probability may be conserved across species, the time scale varies considerably.

The time-dependent increase in growth initiation probability (i.e., Weibull shape >1) may reflect the time dependency of the underlying cellular processes. In the single-cell observations of this study, under low Δ*V_t_* conditions, area at division (𝐴𝑑), generation time (𝑇), and area elongation until division (Δ𝐴) increased in PY1 without CFS compared with the condition with CFS (Fig. S17). These observations indicate that although the cells elongated sufficiently and adequate time had elapsed, cell division was delayed, suggesting that progression toward cell division may be transiently impaired in PY1 in the absence of CFS. Bacterial cell division has been suggested to be closely coordinated with DNA replication initiation and Z-ring assembly [53, 54]. One possible explanation is that the time-dependent increase in growth initiation probability in PY1 reflects the time required for the accumulation of factors involved in these processes, such as DnaA and FtsZ, which may arise from the stochastic nature of transcription and translation [55]. However, the molecular mechanism underlying the time-dependent increase in growth initiation probability remains unclear, and further studies are needed.

Despite these insights, this study has several limitations. First, in the simulation model, stochastic population establishment of PY1 is reproduced under the assumption that Δ*V_t_*-independent growth initiation occurs independently across individual cells. However, growth initiation of individual PY1 cells under extremely low Δ*V_t_* without CFS has not been directly observed (Fig. 2). Thus, it remains unclear whether stochastic population establishment originates from the preceding growth of a single pioneer cell or synchronized growth initiation across the population. Second, the signaling molecules in CFS have not been identified, and the relationship between nitrite and these molecules remains unclear. Therefore, the molecular mechanisms underlying Δ*V_t_*-dependent growth modulation remain to be elucidated, and the representation of the CFS effect as Δ*V_t_* in the simulation has not been directly validated. Addressing these limitations will provide insights into the detailed processes underlying population establishment and the molecular mechanisms that regulate growth in environmental bacteria.

In this study, we demonstrated that combining the Δ*V_t_*-dependent cell growth behavior revealed by single-cell observations with Δ*V_t_*-independent stochastic growth initiation following a Weibull distribution can reproduce both deterministic and stochastic population establishment dynamics in PY1. Stochastic population establishment occurred even in the presence of catalase, whereas the addition of its own CFS induced synchronized population establishment, indicating that PY1 may modulate its growth behavior in response to signaling molecules other than catalase. Furthermore, stochastic population establishment was approximated by a Weibull distribution, a time-dependent stochastic process, implying that time-dependent cellular processes may govern this phenomenon. Notably, in response to CFS, PY1 exhibited generation times and growth rates at the individual cell level comparable to those of the model strain *N. europaea,* thereby leading to rapid and synchronized population establishment. These results indicate that even bacteria previously considered slow-growing and rare can substantially alter their population establishment dynamics depending on environmental conditions. Taken together, our findings suggest that niche differentiation in natural environments may be shaped not only by physicochemical factors, such as temperature, pH, salinity, and substrate concentration, but also by the availability of signaling molecules. We propose that environmentally responsive changes in population establishment dynamics may contribute to sudden blooms of rare taxa in microbial communities.

## Acknowledgments

We gratefully thank John D. McKinney for kindly providing the PDMS microfluidic chips. We thank Rino Isshiki for valuable advice regarding microfluidic device fabrication. We thank Takafumi Inoue for helpful advice regarding the microscopy setup and control. Claude 4.6 and ChatGPT-5 were used to assist with English language editing and translation. The authors have verified the accuracy of the final text.

## Author contributions

All the authors contributed to the conception and design of this study. S.I. performed the experiments and analyses. S.I. wrote the first draft of the manuscript. All authors contributed to manuscript editing.

## Supplementary material

Supplementary material is available at bioRxiv online.

It includes supplementary methods, figures (Figures. S1–S17), and movie legends (Movies S1–S4).

## Data and code availability

All codes for data analysis and simulations can be found in the following GitHub repository: https://github.com/shmy-04/aob-stochastic-growth

