## Supplementary Information for "Environment-responsive individual cell growth behavior shapes stochastic and deterministic population establishment in ammonia-oxidizing bacteria"

**This PDF includes;**
Supplementary methods
Supplementary figures (Figures S1–S17)
Legends for Movies S1–S4

**Other supplementary materials for this manuscript include the following:**
Movie S1 to S4

### Supplementary methods

#### Detailed culture conditions

The inorganic medium used in this study was prepared as previously described and supplemented with NaCl (116 mg $L^{-1}$), $MgSO_4 \cdot 7H_2O$ (40 mg $L^{-1}$), $CaCl_2 \cdot 2H_2O$ (73 mg $L^{-1}$), KCl (38 mg $L^{-1}$), $KH_2PO_4$ (34 mg $L^{-1}$), $FeCl_2$ (2 mg $L^{-1}$), EDTA (4.3 mg $L^{-1}$), $MnCl_2 \cdot 4H_2O$ (0.1 mg $L^{-1}$), $CoCl_2 \cdot 6H_2O$ (0.024 mg $L^{-1}$), $NiCl_2 \cdot 6H_2O$ (0.024 mg $L^{-1}$), $CuCl_2 \cdot 2H_2O$ (0.017 mg $L^{-1}$), $ZnCl_2$ (0.068 mg $L^{-1}$), $Na_2WO_4 \cdot 2H_2O$ (0.033 mg $L^{-1}$), $Na_2MoO_4$ (0.024 mg $L^{-1}$), and $H_3BO_3$ (0.062 mg $L^{-1}$) [1, 2]. The pH of the medium was adjusted to approximately 8.0 using 5 M NaOH, and phenol red was added as a pH indicator.

For strain revival, frozen stocks prepared from 1 mL aliquots of exponential phase cultures were inoculated into 5 mL of medium in glass test tubes (outer diameter 1.75 cm, length 13 cm).

If a decrease in pH was observed during cultivation, the pH was adjusted to approximately 8.0 using 5 M NaOH. This adjustment was performed 1–2 times after subculturing. Cultures were scaled up by subculturing at a ratio of 1–10% (v/v). To minimize the accumulation of DNA mutations, the number of passages after strain revival was limited to five. All experiments were conducted using cultures that had undergone pH adjustment after subculturing and showed a subsequent decrease in pH the following day.

#### Fabrication of microfluidics

In this study, a microfluidic device was constructed as described previously, in which cells were immobilized between a processed cover glass and a cellulose membrane [3].

Cover glass processing was performed as follows. First, circular cover glasses (25-mm in diameter; Thermo Fisher Scientific, USA) were plasma-cleaned using a PR500 plasma cleaner (Yamato Scientific, Japan), and the negative photoresist SU-8 2000.5 (Kayaku Advanced Materials, USA) was spin-coated to a thickness of 750 nm. After soft baking at 95°C for 60 s, UV irradiation of 105 mJ/$cm^2$ was applied using a mask aligner MA6 (SUSS MicroTec, Germany) with a custom-made photomask (Toyo Precision Parts Mfg., Japan). Post-exposure bake was then performed at 95°C for 90 s, followed by development in SU-8 developer (Kayaku Advanced Materials, USA) for 60 s. The cover glasses were subsequently rinsed with IPA for 10 s.

Cellulose membranes with a MWCO of 12–14 kDa (Repligen, USA) were soaked in Milli-Q water for 2–3 h and then cut into discs approximately 25 mm in diameter. The cut membranes were washed again in Milli-Q water for 1 h, followed by an overnight immersion in methanol for sterilization. After sterilization, the membranes were sandwiched between autoclaved filter paper and dried in a vacuum chamber for at least 48 h.

#### Detailed microscope configurations

Time-lapse images were acquired using an inverted microscope IX81-ZDC (Evident, Japan) equipped with a CoolSNAP $_{HQ2}$ Monochrome camera (6.45 μm pixel size; Photometrics, USA) and a 100×/1.40 oil UPlanSApo objective (Evident, Japan), resulting in an effective pixel size of 64.5 nm. Approximately 10 fields of view (FOV) were imaged at each time point using a motorized stage MPT-AS01-FV (SIGMAKOKI, Japan). The Z-axis drift during imaging was corrected using the ZDC function. During the observation, the device was placed on a stage heater INUG2-ONI (Tokai Hit, Japan) to maintain a constant temperature. Bright-field images were acquired every 10 min with halogen lamp illumination for approximately 5 s per acquisition. All microscope components were controlled using a Ti Workbench [4]. To minimize the Z-axis drift caused by temperature fluctuations, observations were initiated 3 h after the microscope was turned on.

#### Detailed Image processing and cell tracking procedures

Image processing was performed using a sequential pipeline. First, uneven illumination was corrected, and contrast was enhanced using Fiji [5]. Positional drift in the XY plane was corrected using StackReg and BigWarp [6, 7]. Next, a StarDist model was trained based on previously described procedures to generate a custom StarDist model [8–10]. Cell segmentation was performed using the trained StarDist model in combination with StarDist-OPP [11]. Segmented cells were then separated using the Adjustable Watershed algorithm (tolerance = 3.0) [12]. Individual cell tracking was subsequently performed using custom Python code. Finally, cell lineages were constructed using TrackMate [13]. All cell lineages were manually inspected and corrected during TrackMate-based lineage construction. Cells that were temporarily lost were manually reconnected to their lineages if they were detected again within 3 frames.

To improve analysis throughput, images were analyzed every 10 frames (every 100 min) for PY1 and every 5 frames (every 50 min) for *N. europaea*. When the field of view (FOV) was unsuitable for analysis owing to temporary Z-axis defocus, the affected frame was replaced with an image from the immediately preceding or following frame.

The following cells were excluded from subsequent analyses:

(1) Cells without a traceable ancestor (e.g., cells that entered the FOV during observation)

(2) Cells located at the edge of the image, for which the cell area could not be accurately quantified

(3) Cells that became inseparable from neighboring cells and could no longer be tracked as individual cells during image analysis

(4) Cells for which no cell division was confirmed during the observation period; therefore, the generation time could not be defined

#### Estimation of biomass production $\Delta V_t$

Biomass production ($\Delta V_t$) was defined as:

$$\Delta V_t = \frac{V_{\mathrm{biomass}}(t) \times (\exp{(\alpha_{\mathrm{mean}}(t) \times \Delta t)} - 1)}{V_{\mathrm{medium}}}$$

where $V_{\mathrm{biomass}}(t)$ is the total cell volume within the FOV at time $t$, $\alpha_{\mathrm{mean}}(t)$ is the mean elongation rate of cells within the FOV at time $t$, $\Delta t$ is the time interval used for analysis (PY1: 100 min; *N. europaea*: 50 min), and $V_{\mathrm{medium}}$ is the volume of medium flowing into the FOV during $\Delta t$.

$V_{\mathrm{biomass}}(t)$ was calculated by multiplying the total cell area within the FOV at time $t$ ($A_{\mathrm{total}}(t)$) by the height of the culture chamber constructed on the cover glass (750 nm). For $A_{\mathrm{total}}(t)$, an estimated value was obtained by fitting the time series of the total measured cell area within the FOV to an exponential growth curve ($A_{\mathrm{total}}(t) = A_{\mathrm{total}}(0) \times e^{\alpha t}$). In one FOV of *N. europaea*, substantial cell outflow from the FOV was observed as cell number increased. Therefore, only data up to the time point at which the measured cell area reached its maximum were used for the FOV.

$\alpha_{\mathrm{mean}}(t)$ was calculated as the mean elongation rate ($\alpha$) at time $t$ obtained from individual cell tracking. The calculation of $\alpha_{\mathrm{mean}}(t)$ was restricted to time points at which the number of tracked cells was $\geq$10.

$V_{\mathrm{medium}}$ was calculated by multiplying the volume of medium flowing into the device during the time interval $\Delta t$ by the ratio of the area of the observed FOV to the total area of the device. The total area of the device was estimated to be approximately $1.47 \times 10^8$ µm$^2$ based on the design specifications reported previously [3]. For example, in the case of PY1, the volume of medium flowing into the device during $\Delta t$ was estimated to be 800 µL (8 µL min$^{-1}$ × 100 min). Assuming an observed FOV of 1392 × 1040 pixels, the FOV area is $1392 \times 1040 \times 0.0645^2 = 6.02 \times 10^3$ µm$^2$.

Consequently, $V_{\mathrm{medium}}$ = 800 μL × (6.02 × $10^3$ $\mu m^2$ / 1.47 × $10^8$ $\mu m^2$) = 3.28 × $10^{-2}$ μL.

#### Simulation model for microfluidics

The parameters required for the simulation ($A_0$, $\alpha_0$, $T_0$, $A_{\mathrm{min,div}}$, $DR$, $\alpha$, $T$ and $A_{\mathrm{max}}$) were estimated from single-cell observations of PY1 in the absence of CFS using kernel density estimation (KDE; $\alpha_0$ and $T_0$), random forest prediction ($A_0$), and regression models ($A_{\mathrm{min,div}}$, $DR$, $\alpha$, $T$ and $A_{\mathrm{max}}$) (Fig. S9). Outliers were removed using a boxplot criterion (1.5 × IQR) based on the elongation rate to reduce the influence of extreme values with low observation counts and to enable robust parameter estimation and regression.

During simulation, $\Delta V_t$ was calculated at each time step, and the simulation was terminated when $\Delta V_t$ reached 3.0 × $10^6$ $\mu m^3$ $mL^{-1}$. Because the microfluidic device operated as a continuous culture system, the accumulation of metabolites was assumed to be negligible. The simulations were repeated the same number of times as the number of FOV analyzed (biological replicates, n = 3; 3 FOV per replicate). The initial number of cells in each simulation was set to match the number of cells successfully tracked in the corresponding FOV (replicate 1: 45, 28, 42; replicate 2: 68, 76, 69; replicate 3: 57, 56, 40 cells). The simulation code to reproduce cell behavior in a microfluidic device was developed based on a previously described study [14].

#### Evaluation of simulation accuracy using JSD

To evaluate the agreement between experimental and simulation results within the microfluidic device, the Jensen–Shannon divergence (JSD) between the experimental and simulation data distributions was calculated.

First, both the experimental and simulation data were divided into 10 bins based on $\Delta V_t$ at cell birth. Random subsampling was performed on the simulation data to match the sample sizes of the experimental and simulation data within each bin.

Next, for each individual cell growth parameter, the distributions were estimated for each bin using KDE. The JSD between the experimental and simulated data distributions was calculated for each bin.

JSD calculation was repeated 30 times to account for the stochasticity introduced by subsampling, and the mean value was used as the evaluation metric. The five individual cell growth parameters ($T$, $\alpha$, $\Delta A$, $Ab$, and $Ad$) were evaluated based on a previous study [15].

#### Simulation model for batch culture

The simulation of PY1 cell growth behavior in batch culture was performed in the same manner as in the microfluidic device simulation. However, in the batch culture simulation, metabolites were assumed to accumulate within the system, including $\Delta V_t$ and nitrite.

At the start of the simulation, the initial cell area $A_0$ was drawn from a uniform distribution between $A_{\mathrm{min,div}}/2$ and $A_{\mathrm{min,div}}$. The initial elongation rate $\alpha_0$ and initial generation time $T_0$ were calculated using the same regression model applied to cells from generation 1 onward, based on $A_0$ and initial $\Delta V_t$ at the start of cultivation. The initial number of cells was drawn from a Poisson distribution with the expected cell count corresponding to the inoculation densities ($10^1$, $10^3$, $10^5$ cells $mL^{-1}$) as the mean ($\lambda$).

Nitrite production was calculated as the accumulated $\Delta V_t$ divide by the steady-state cell birth volume and multiplied by the growth yield of PY1 (33.5 cells $pmol^{-1}$) [2]. The steady-state cell birth volume was estimated by multiplying the steady-state cell area at birth observed within the microfluidic device ($A_{\mathrm{min,div}}/2$) by the height of the culture chamber on a cover glass (750 nm).

To represent the decrease in growth rate due to nitrite accumulation, the inhibition constant $K_i$ for nitrite was introduced. The effective cell elongation rate was limited according to a noncompetitive inhibition model based on the nitrite concentration $[NO_2^-(t)]$:

$$\alpha_{\mathrm{eff}} = \alpha \times \frac{1.0}{\left\{1.0 + \left(\frac{[NO_2^-(t)]}{K_i}\right)\right\}}$$

Note that $K_i$ in this study does not represent the inhibitory effect of nitrite itself. Instead, it was treated as an apparent inhibition constant encompassing unmodeled factors in the culture system, such as free nitrous acid and free ammonia concentrations associated with pH fluctuations. To reproduce the experimentally observed nitrite production dynamics in the presence of CFS, $K_i$ was set to 100 µM (Fig. S11).

#### Fitting the initiation probability of $\Delta V_t$–independent cell growth

To estimate the probability of $\Delta V_t$-independent growth initiation, the probability of population establishment arising from such cells was modeled based on a previously described framework, with modifications (Fig. S14) [16]. In this model, $F(t)$ was defined as the cumulative probability that an individual cell initiates growth by time $t$. Given the initial cell number ($N_{\mathrm{cells}}$), the expected number of populations formed by at least one growth-initiating cell ($N(t)$) can be expressed as:

$$N(t) = N_{\infty} \times \{1 - (1 - F(t))^{N_{\mathrm{cells}}}\}$$

where $N_{\infty}$ denotes the theoretical maximum number of detectable population establishments. For $F(t)$, two distributions were assumed: an exponential distribution based on a Poisson process, and a Weibull distribution in which the probability increases with the duration of the lag phase.

For the Poisson process, the following equation was used:

$$F(t) = 1 - e^{-\lambda(t - t_r - lag)}$$

For the Weibull distribution, the following equation was used:

$$F(t) = 1 - exp\left(-\frac{t - t_r{}^{k}}{\tau}\right)$$

where $t_r$ is the time required for an individual cell to produce the threshold amount of nitrite for detection (0.25 mM) after growth initiation, and $lag$ denotes the minimum lag time before a $\Delta V_t$-independent growth-initiating cell emerges. The value of $t_r$ was set to 289.92 h, corresponding to the time required for PY1 to produce 0.25 mM nitrite in simulations assuming its ideal elongation rate and generation time. The ideal elongation rate and generation time of PY1 were set to the values derived from single-cell observations in the absence of CFS (Fig. S9): a generation time of approximately 6.14 h and an elongation rate of approximately 0.064 $h^{-1}$. Furthermore, $\lambda$ represents the rate of growth initiation in the Poisson process, whereas $k$ and $\tau$ represent the shape and scale parameters of the Weibull distribution, respectively. These parameters define the probability of $\Delta V_t$-independent growth initiation.

#### Simulation model for batch culture with Weibull hazard

In the batch culture simulation incorporating $\Delta V_t$-independent growth initiation, the Weibull hazard function $h(t)$ was used to model the emergence of cells initiating $\Delta V_t$-independent growth. The Weibull hazard function was calculated as follows:

$$h(t) = \frac{k}{\tau} \times \left(\frac{t}{\tau}\right)^{k-1}$$

where $t$ represents cell age, corresponding to the duration of the lag phase. The probability of a growth initiation event occurring within the simulation time step $dt$ was calculated as follows:

$$p(t) = 1 - \exp\left(-h(t) \times dt\right)$$

At each simulation step, $p(t)$ was calculated, and a random number between 0 and 1 was generated for each cell that had not yet initiated $\Delta V_t$-independent growth. If the generated random number was smaller than $p(t)$, the cell was defined as initiating $\Delta V_t$-independent growth. Cells initiating $\Delta V_t$-independent growth were assumed to elongate and divide according to values derived from single-cell observations of PY1 in the absence of CFS (generation time $\approx$ 6.14 h and elongation rate $\approx$ 0.064 $h^{-1}$).

### Supplementary figures

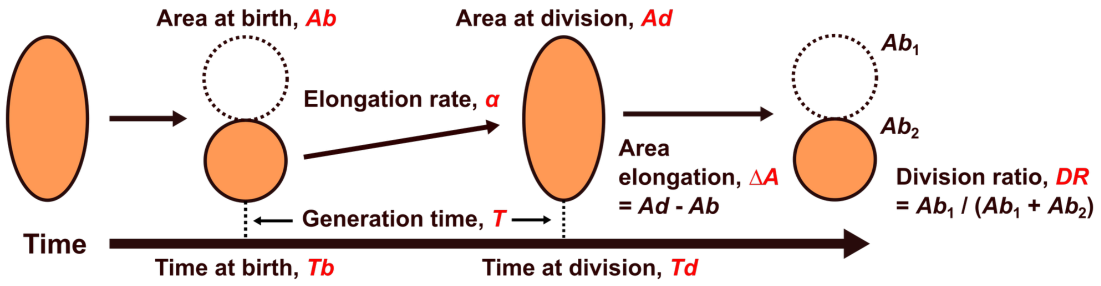


**Figure S1. Definition of eight parameters extracted from single-cell observations.** The following parameters were used to characterize the individual cell growth behavior of AOB: time of birth ($Tb$), time of division ($Td$), generation time ($T$), area at birth ($Ab$), area at division ($Ad$), elongation rate ($\alpha$), cell area elongation until division ($\Delta A$), and division ratio ($DR$).

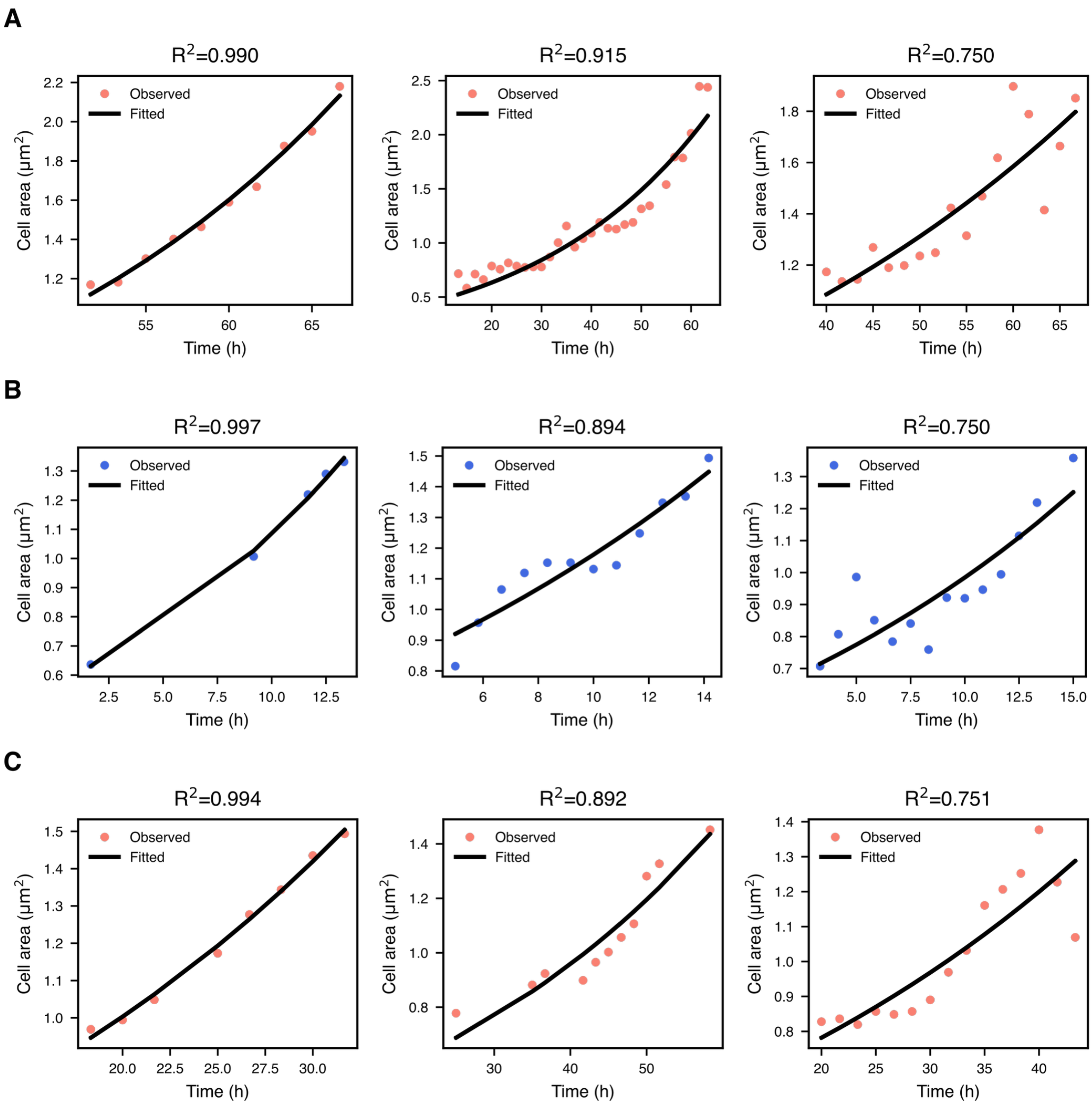


**Figure S2. Exponential growth fitting of individual cell area trajectories.** Area trajectories of individual cells extracted from single-cell observations were fitted to an exponential growth curve ($A(t) = Ab \times e^{\alpha t}$). Cells with a coefficient of determination ($R^2$) >0.75 were sampled at regular intervals of $R^2$, and the area trajectories together with the fitting curves were plotted. Filled circles represent experimental data, and black lines represent the fitted curves. Cell area was calculated by multiplying the number of pixels per cell by the area per pixel (0.0645 × 0.0645 μm$^2$). Cells with $R^2$ >0.75 were used in subsequent analyses, and $Ab$, $Ad$, and $\alpha$ were calculated from the fitting results for each cell. (A) PY1 without CFS. (B) *N. europaea* without CFS. (C) PY1 with CFS.

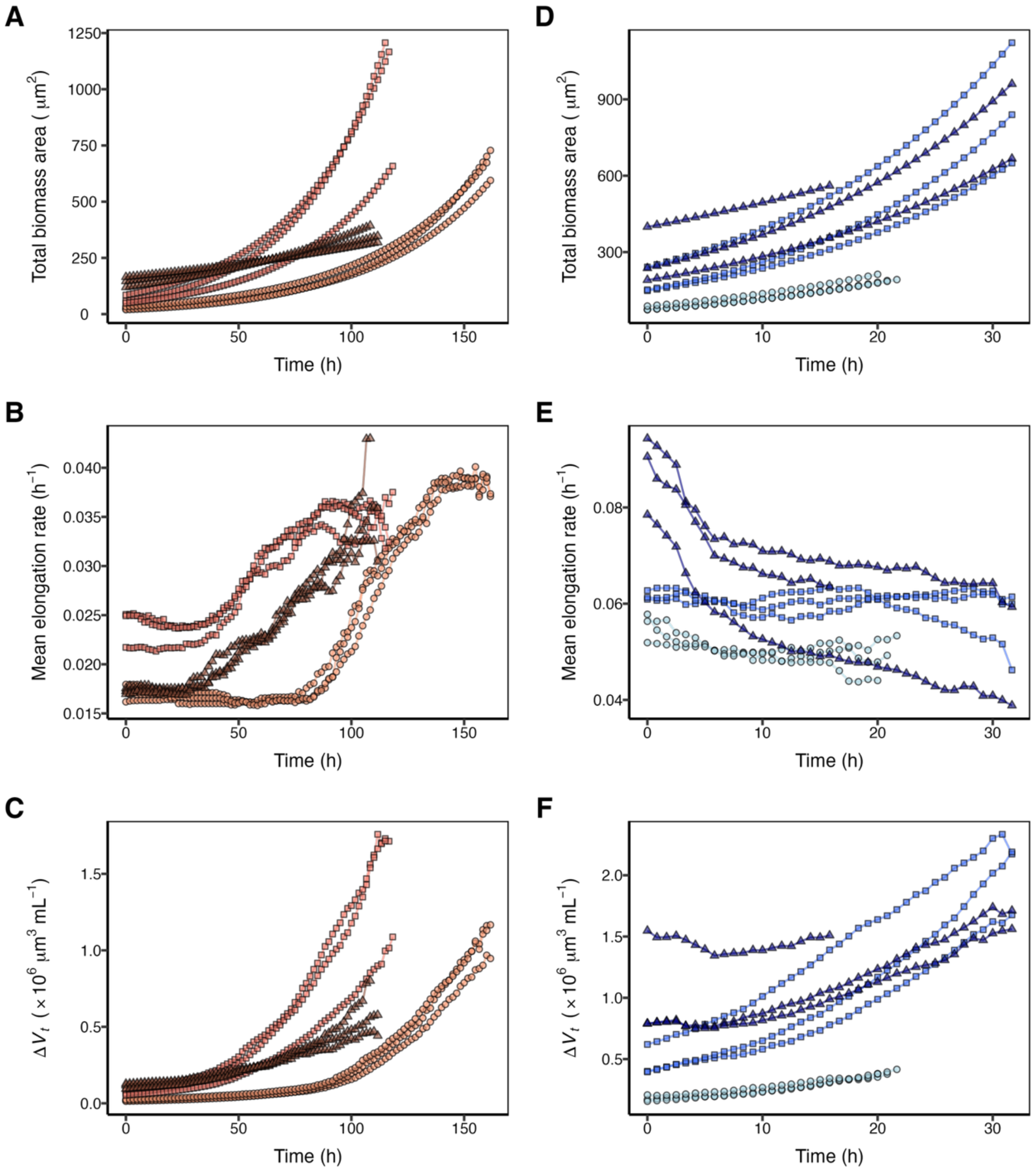


**Figure S3. Temporal changes in biomass, mean elongation rate, and biomass production ($\Delta V_t$) within each field of view (FOV) in two Nitrosomonas strains.** (A–C) PY1. (D–F) *N. europaea*. (A, D) Total biomass area within the FOV. (B, E) Mean cell elongation rate within the FOV. (C, F) $\Delta V_t$ within the FOV. Symbols and colors indicate biological replicates (n = 3; 3 FOVs per replicate).

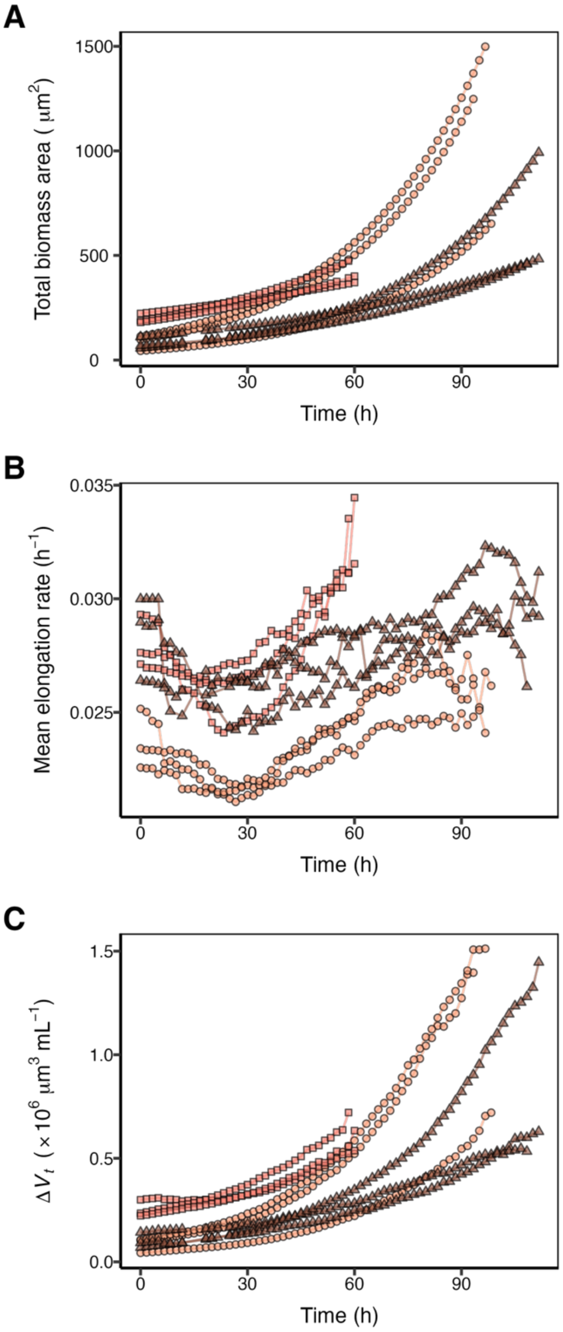


**Figure S4. Temporal changes in biomass, mean elongation rate, and biomass production ($\Delta V_t$) within each field of view (FOV) for PY1 with CFS.** (A) Total biomass area within the FOV. (B) Mean cell elongation rate within the FOV. (C) $\Delta V_t$ within the FOV. Symbols and colors indicate biological replicates (n = 3; 3 FOVs per replicate).

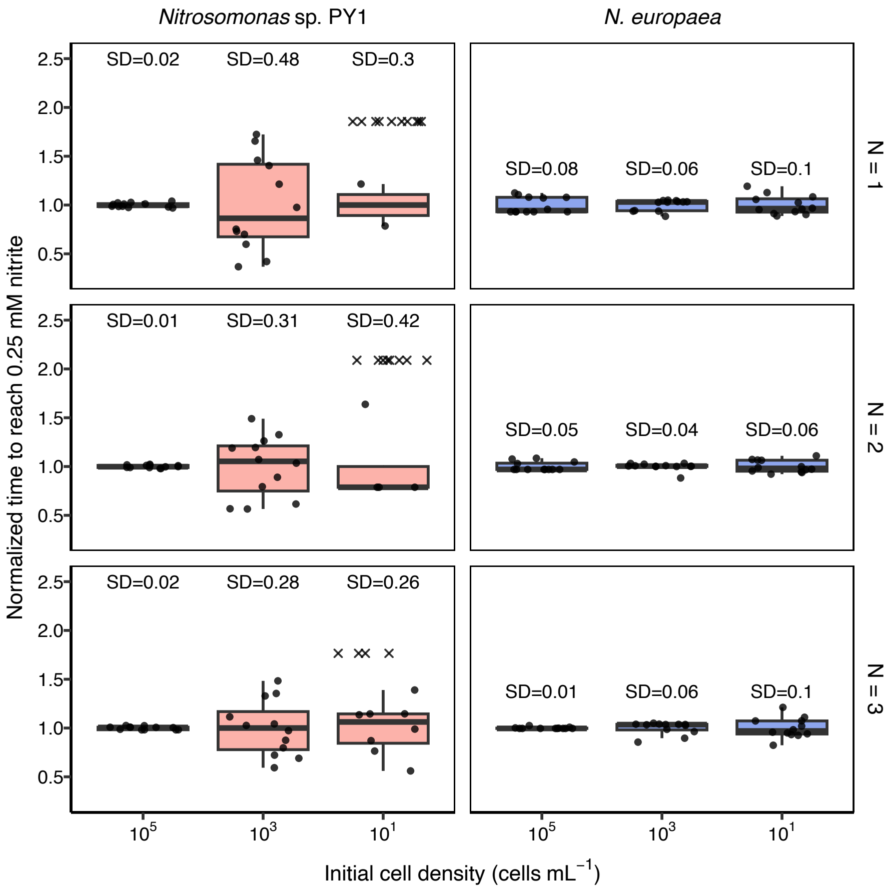


**Figure S5. Heterogeneity in population establishment timing in two *Nitrosomonas* strains.** Variability in nitrite detection time when each strain was inoculated into 48-well plates at $10^5$, $10^3$, and $10^1$ cells $mL^{-1}$ in the absence of CFS. To eliminate differences in time scale due to strain-specific growth rates, values normalized by the mean nitrite detection time under each condition are plotted. Experiments were performed in biological replicates (n = 3). Wells in which nitrite concentration did not reach 0.25 mM during the observation period are indicated by crosses (×). Data from replicate 1 are shown in the main text (Fig. 4B).

Two *Nitrosomonas* strains were inoculated into 48-well plates at $10^5$, $10^3$, and $10^1$ cells $mL^{-1}$ in the absence of CFS, and the time at which nitrite concentration exceeded 0.25 mM was defined as the nitrite detection time. The detection times were normalized to the mean detection time under each culture condition before plotting.

In PY1, particularly at an initial cell density of $10^3$ cells $mL^{-1}$, the normalized nitrite detection time showed substantial variability, with a standard deviation approximately 7-fold larger than that in *N. europaea*. At the lowest initial density ($10^1$ cells $mL^{-1}$), several PY1 wells did not reach detectable nitrite levels within the observation period (269 days).

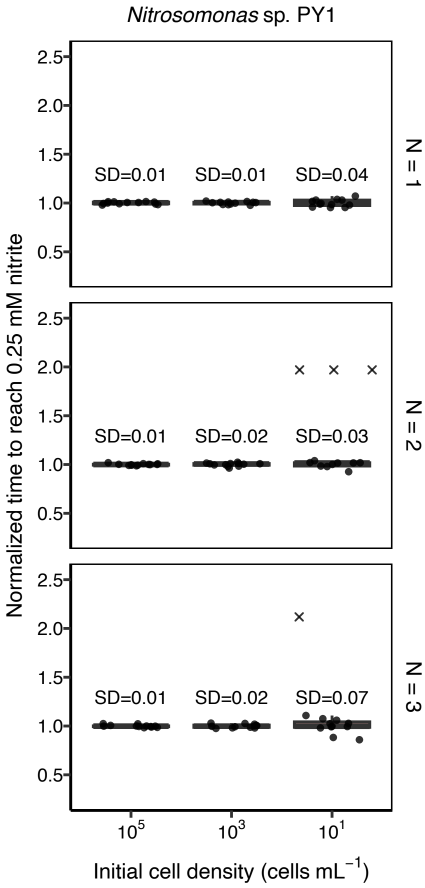


**Figure S6. Heterogeneity in population establishment timing of PY1 in the presence of CFS.** Variability in nitrite detection time when PY1 was inoculated into 48-well plates at $10^5$, $10^3$, and $10^1$ cells $mL^{-1}$ in the presence of CFS. Values normalized by the mean nitrite detection time under each condition are plotted. Experiments were performed in biological replicates (n = 3). Wells in which nitrite concentration did not reach 0.25 mM during the observation period are indicated by crosses (×). Data from replicate 1 are shown in the main text (Fig. 4D).

PY1 was inoculated into 48-well plates at $10^5$, $10^3$, and $10^1$ cells $mL^{-1}$ in the presence of CFS, and the time at which nitrite concentration exceeded 0.25 mM was defined as the nitrite detection time. The detection times were normalized to the mean detection time under each culture condition before plotting.

At an initial cell density of $10^3$ cells $mL^{-1}$, the variability in the normalized nitrite detection time was substantially reduced, with a standard deviation approximately 22-fold smaller than that in the absence of CFS.

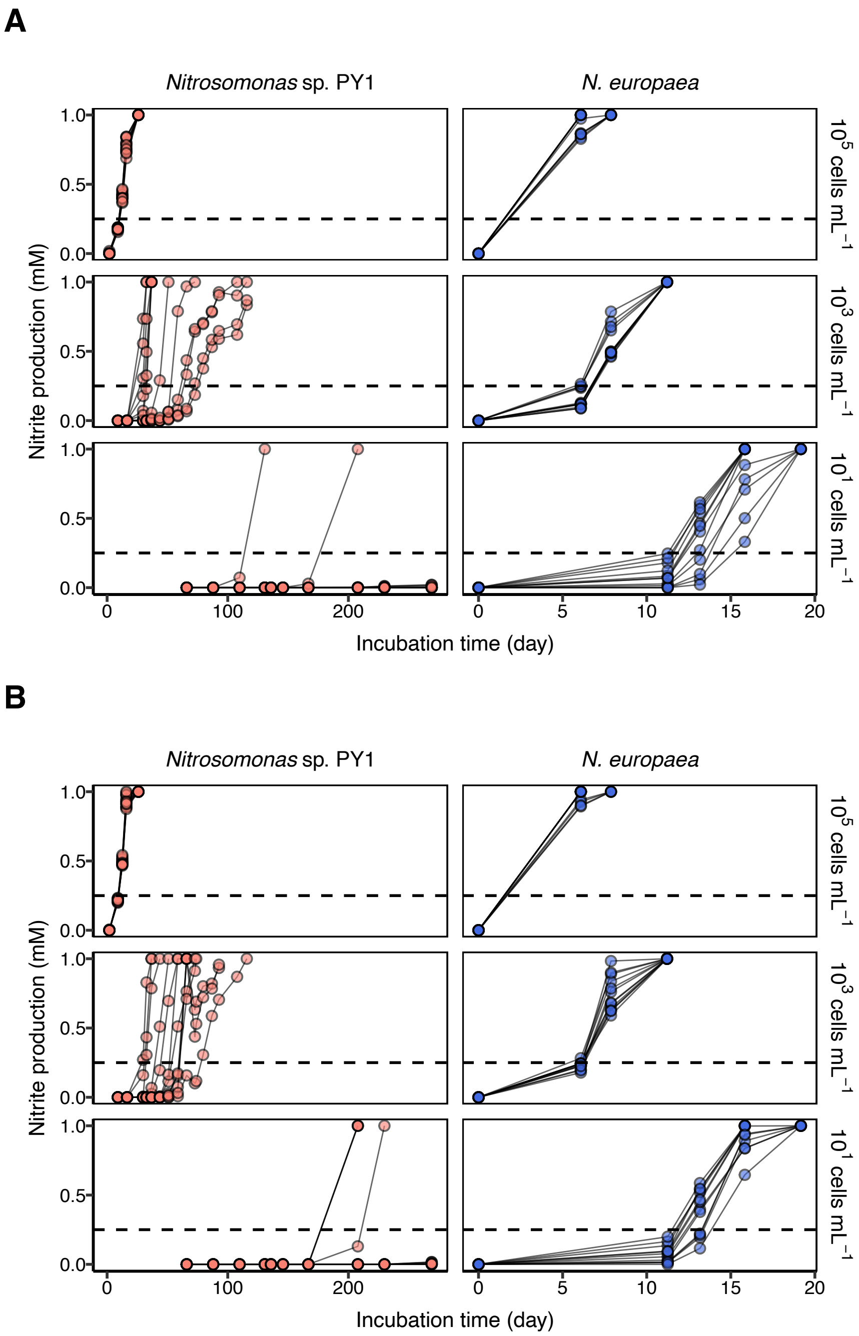

A
*Nitrosomonas* sp. PY1
*N. europaea*
Nitrite production (mM)
1.0
0.5
0.0
$10^5$ cells $mL^{-1}$
$10^3$ cells $mL^{-1}$
$10^1$ cells $mL^{-1}$
0
100
200
0
5
10
15
20
Incubation time (day)
B
*Nitrosomonas* sp. PY1
*N. europaea*
Nitrite production (mM)
1.0
0.5
0.0
$10^5$ cells $mL^{-1}$
$10^3$ cells $mL^{-1}$
$10^1$ cells $mL^{-1}$
0
100
200
0
5
10
15
20
Incubation time (day)

C

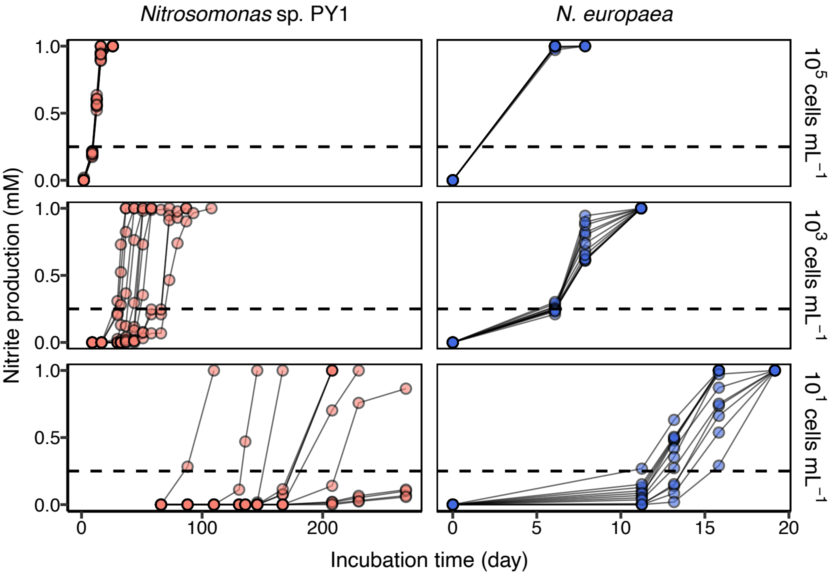


**Figure S7. Time course of nitrite concentration in batch culture experiments for two *Nitrosomonas* strains in the absence of CFS.** Nitrite concentration was monitored after each strain was inoculated into 48-well plates at $10^5$, $10^3$, and $10^1$ cells $mL^{-1}$ in the absence of CFS. (A–C) Data for each biological replicate (n = 3). The black dotted line indicates a nitrite concentration of 0.25 mM. These data were used to generate Figs. 4A, 4B, and S5.

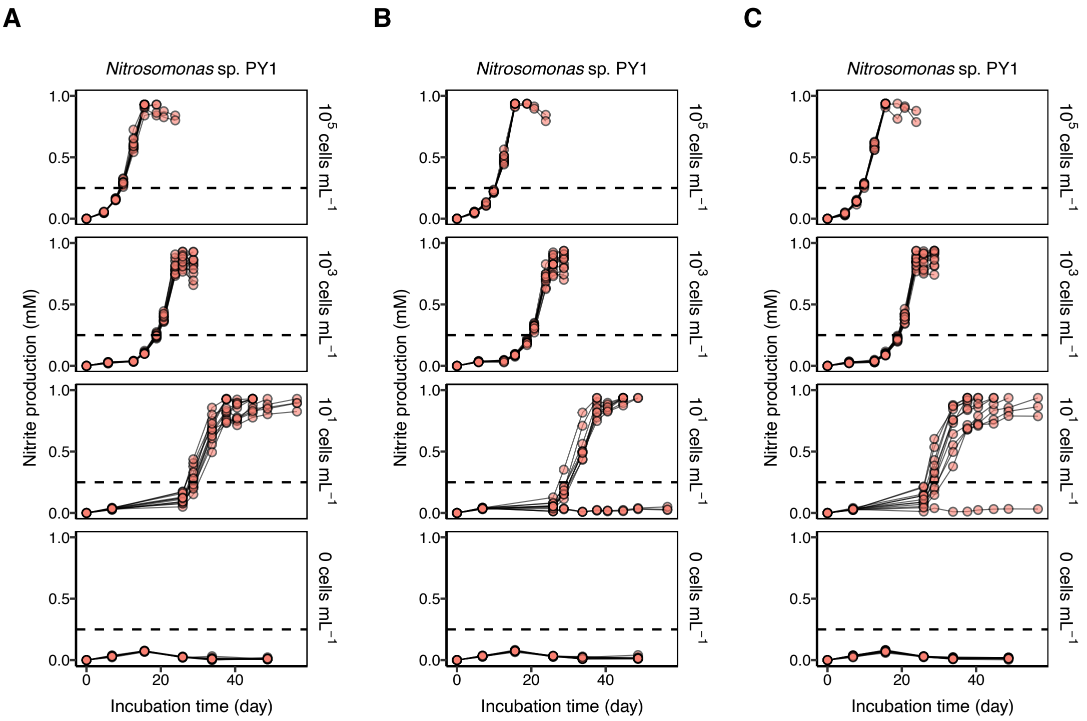


**Figure S8. Time course of nitrite concentration in batch culture experiments for PY1 in the presence of CFS.** Nitrite concentration was monitored after PY1 was inoculated into 48-well plates at $10^5$, $10^3$, $10^1$, and a no-inoculum control (0 cells $mL^{-1}$) in the presence of CFS. (A–C) Data for each biological replicate (n = 3). The black dotted line indicates a nitrite concentration of 0.25 mM. These data were used to generate Figs. 4C, 4D, and S6.

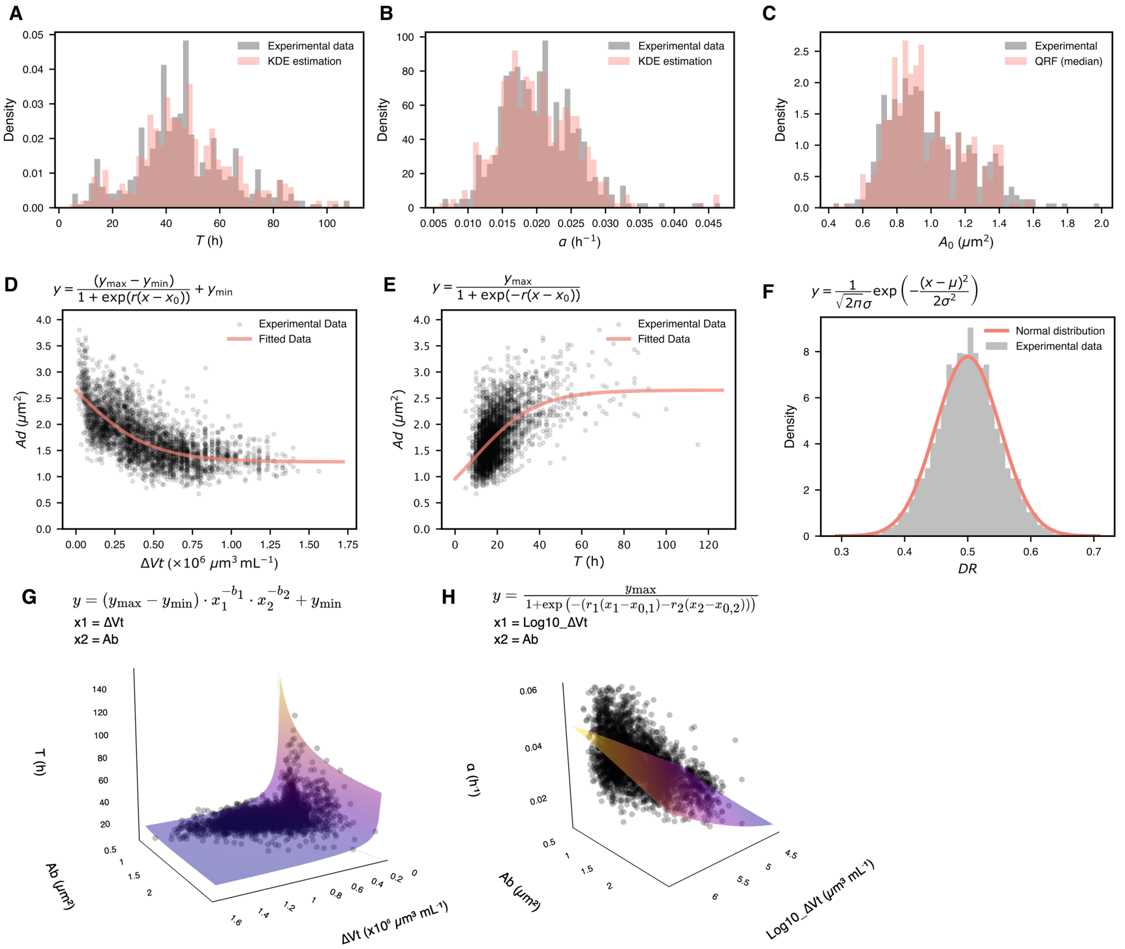


**Figure S9. Models for reproducing individual cell growth parameters within the microfluidic device for PY1 in the absence of CFS.** Individual cell growth parameters were extracted from the analysis of single-cell observations performed in the absence of CFS. Models reproducing the distribution of each parameter were constructed using kernel density estimation (A, B), random forest (C), and parametric models (D–H). (A–C) Models for initial cell growth parameters. Histograms and reproduced distributions for initial generation time ($T_0$), initial elongation rate ($\alpha_0$), and initial cell area ($A_0$) of generation 0 cells. $T_0$ and $\alpha_0$ were reproduced using kernel density estimation (KDE), and $A_0$ was reproduced using quantile random forest (QRF) with $T_0$ and $\alpha_0$ as input variables. (D-F) Models for cell size and division parameters. The minimum cell area required for division ($A_{\mathrm{min,div}}$) was estimated using a logistic decay model with biomass production ($\Delta V_t$) as the explanatory variable. The asymptotic value ($y_{\mathrm{min}}$) was defined as $A_{\mathrm{min,div}}$ (D). The maximum cell area ($A_{\mathrm{max}}$) was estimated using a logistic model with generation time ($T$) as the explanatory variable. In the batch culture simulation, the asymptotic value of the logistic model ($y_{max}$) was used as $A_{\mathrm{max}}$ (E). The division ratio ($DR$) was assumed to follow a normal distribution parameterized by the mean and standard deviation of the experimental data (F). (G-H) Regression models describing growth dynamics of cells from generation 1 onward. Generation time ($T$) was modeled using $\Delta V_t$ and cell area at birth ($Ab$) with an exponential decay model (G). Elongation rate ($\alpha$) was modeled using biomass production ($\Delta V_t$) and $Ab$ with a logistic decay model (H). These regression models were used to reproduce $T$ and $\alpha$ in all simulations. The asymptotic values at high $\Delta V_t$ ($T = 6.14$ h, $\alpha = 0.064$ $h^{-1}$) were used as ideal growth conditions; see Code Availability for details.

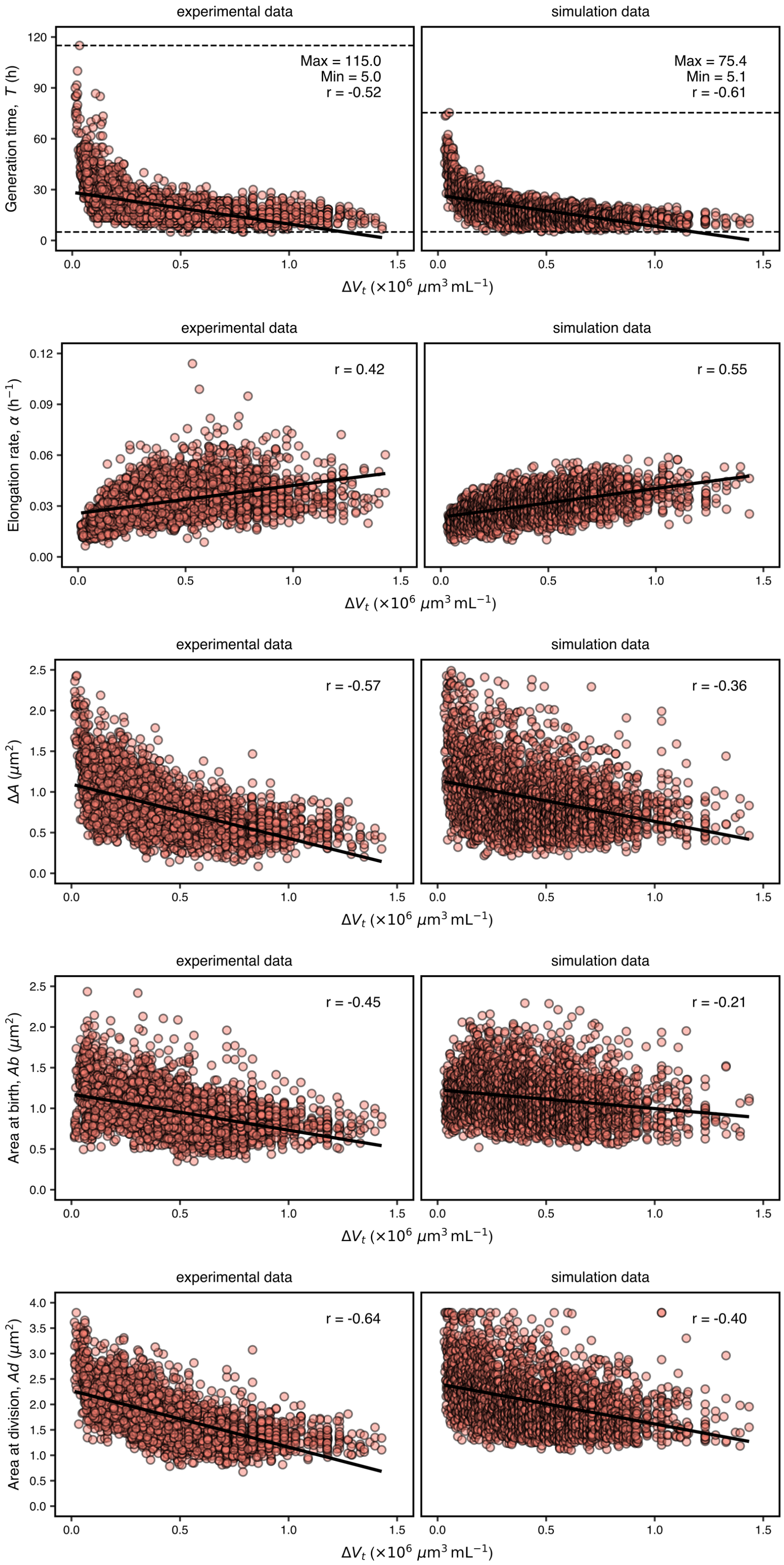
experimental data
simulation data
Generation time, T (h)
Max = 115.0
Min = 5.0
r = -0.52
Max = 75.4
Min = 5.1
r = -0.61
Elongation rate, α (h⁻¹)
r = 0.42
r = 0.55
ΔA (μm²)
r = -0.57
r = -0.36
Area at birth, Ab (μm²)
r = -0.45
r = -0.21
Area at division, Ad (μm²)
r = -0.64
r = -0.40
ΔVt (×10⁶ μm³ mL⁻¹)

**Figure S10. Comparison of experimental and simulated individual cell growth parameters of PY1 within the microfluidic device in the absence of CFS.** Growth behavior of PY1 within the device was simulated using the model described in Fig. 5A. Experimental (left) and simulated (right) individual cell growth parameters ($T$, $\alpha$, $\Delta A$, $Ab$, $Ad$) were plotted as a function of biomass production ($\Delta V_t$). Data were divided into 10 bins based on $\Delta V_t$, and random sampling was performed from the simulation results so that the number of data points in each bin matched that of the experimental data. Linear regression was performed for each plot, and the correlation coefficients are shown.

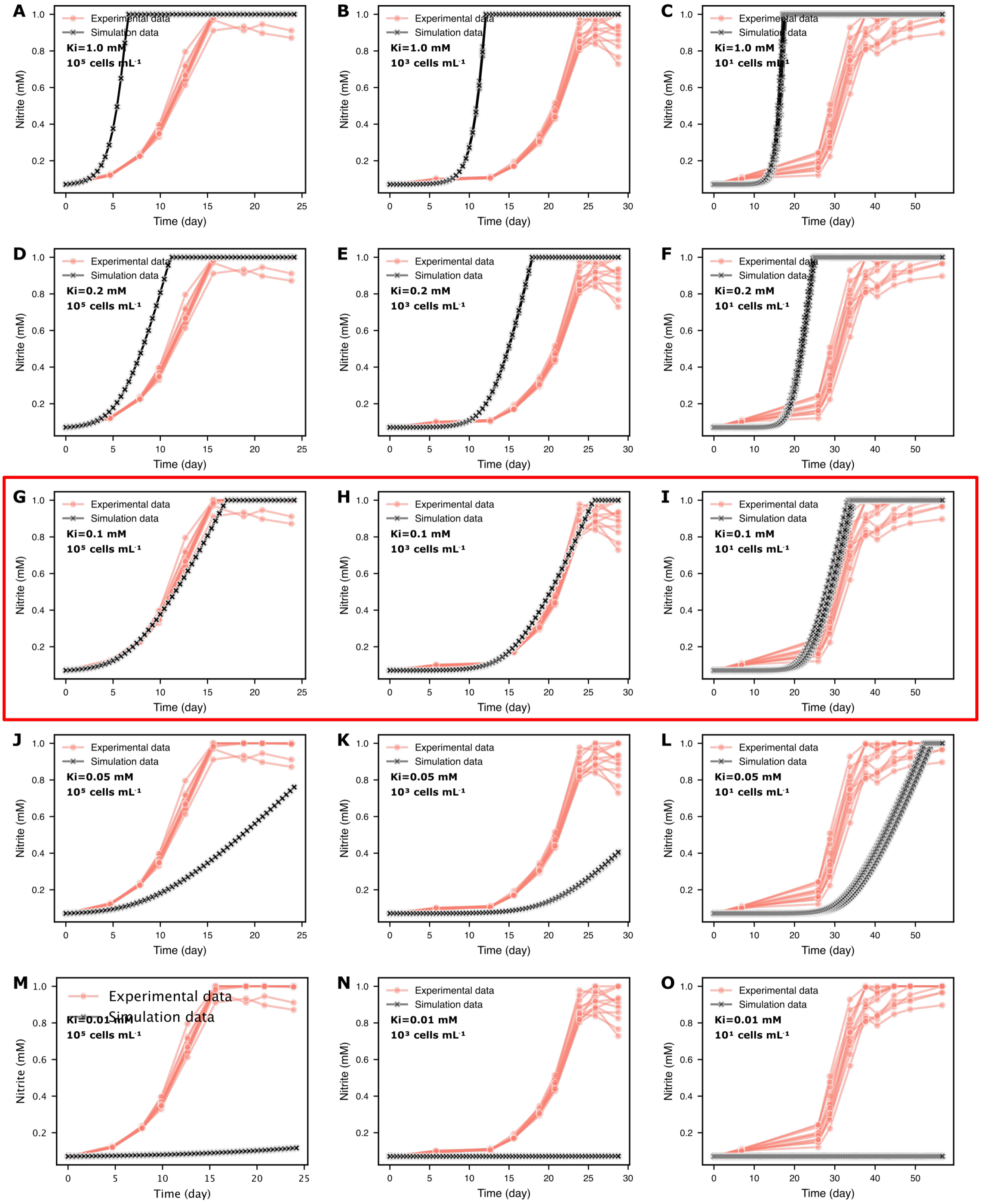


**Figure S11. Evaluation of the apparent inhibition constant ($Ki$) in a simulation model for PY1 in batch culture.** The $Ki$ for nitrite was varied, and growth of PY1 cells under batch culture was reproduced using the same individual-based model (IBM) and parameter settings as in Fig. 6A. Simulated time courses of nitrite concentration derived from the growth simulation were compared with experimental data. Red circles and lines represent experimental data, and black triangles and lines represent simulation results. The initial cell number was stochastically assigned from a Poisson distribution with mean $\lambda$, set to $10^5$, $10^3$, and $10^1$ cells. Experimental data from a representative biological replicate and the corresponding simulation results are shown. $Ki$ was set to 1.0 mM (A–C), 0.2 mM (D–F), 0.1 mM (G–I), 0.05 mM (J–L), and 0.01 mM (M–O). Note that no buffer was added in the batch culture of PY1, resulting in pH fluctuations and associated changes in free ammonia and free nitrous acid concentrations. These changes affect the concentrations of substrates, inhibitors, and metabolites, but are difficult to incorporate into the model. Therefore, $Ki$ was treated as a phenomenological parameter representing unmodeled factors that reduce growth rate with increasing nitrite concentration.

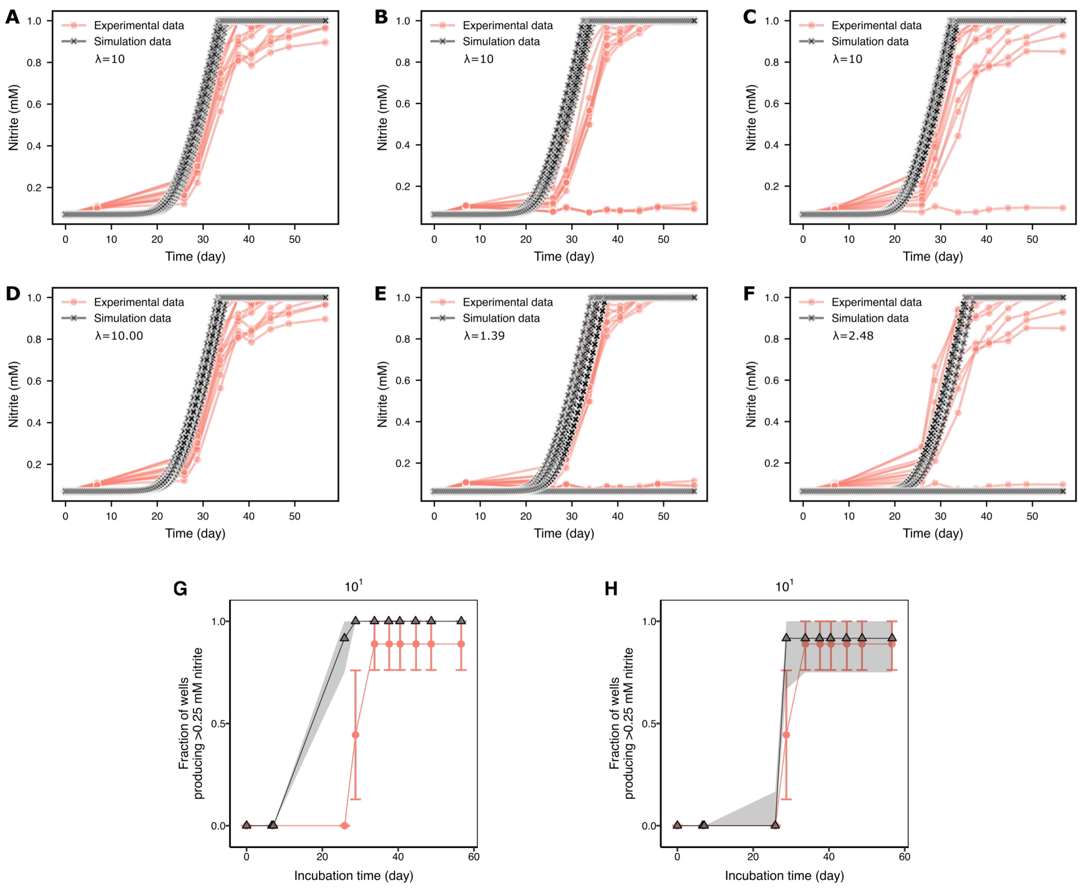


**Figure S12. Simulation results for PY1 in batch culture with varying expected initial cell count ($\lambda$).** Simulations were performed using the same individual-based model (IBM) and parameter settings as in Fig. 6A, while varying $\lambda$. The analysis focused on conditions assuming an expected initial cell density of $10^1$ cells $mL^{-1}$. Red circles and lines represent experimental data, and black triangles and lines represent simulation results. (A–C) Time course of nitrite concentration per well with $\lambda = 10$. (D–F) Time course of nitrite concentration per well under conditions with adjusted $\lambda$. For biological replicates in which nitrite-negative wells were observed, $\lambda$ was estimated as $-ln(p0)$, where $p0$ is the observed proportion of nitrite-negative wells. (G) Simulations were performed 100 times independently with $\lambda = 10$, and the median proportion of nitrite-positive wells at each time point is shown. The 95% prediction interval is indicated by a shaded region. Wells in which nitrite concentration reached $\geq$0.25 mM were defined as nitrite-positive wells. (H) The same analysis as in (G) was performed with $\lambda = 2.48$.

As shown in Fig. 6A, under the condition assuming an expected initial cell density of $10^1$ cells $mL^{-1}$ in the presence of CFS, nitrite detection in the simulation occurred slightly earlier than that in the experiment. Plotting the time course of the nitrite concentration in each well confirmed that the onset of nitrite increase in the simulation preceded that in the experiment by a few days (Fig. S12A–C).

Under low cell density conditions, stochastic variability during inoculation may result in zero-cell wells. Indeed, in the batch culture experiment with an initial cell density of $10^1$ cells $mL^{-1}$, nitrite production was not detected in some wells, even in the presence of CFS (Fig. S8B, C). Assuming a Poisson distribution of cells among wells, the probability of zero-cell wells is given by $p0 = \exp(-\lambda)$. Therefore, $\lambda$ can be estimated as $-ln(p0)$.

Using this approach, $\lambda$ was estimated for the two biological replicates in which nitrite-negative wells were observed ($\lambda = 1.39, 2.48$). Simulations using these estimated $\lambda$ reproduced the appearance of nitrite-negative wells and yielded nitrite production dynamics more consistent with the experimental results (Fig. S12D–F). Furthermore, simulations with $\lambda = 2.48$ more accurately reproduced the dynamics of nitrite-positive well proportions than those with $\lambda = 10$ (Fig. S12G, H).

These results suggest that, although the nominal initial cell density was set to $10^1$ cells $mL^{-1}$, the effective initial cell density was lower. This discrepancy may arise from cell counting errors or inaccuracies during dilution.

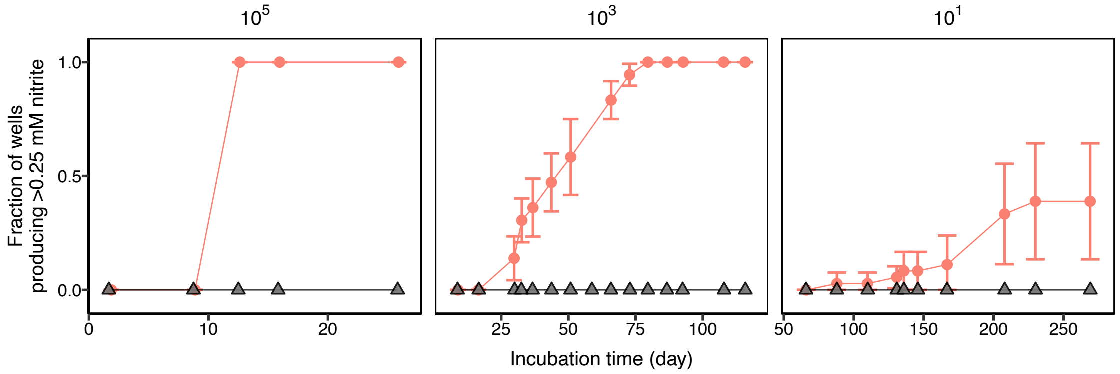


**Figure S13. Simulation of the proportion of nitrite-positive wells for PY1 in batch culture with initial $\Delta V_t = 1$ in the absence of CFS.** Simulations were performed using the same individual-based model (IBM) and parameter settings as in Fig. 6A. Red circles and lines represent experimental data, and black triangles and lines represent simulation results. Simulations were performed 100 times independently, and the median proportion of nitrite-positive wells at each time point is shown. The 95% prediction interval is indicated by a shaded region.

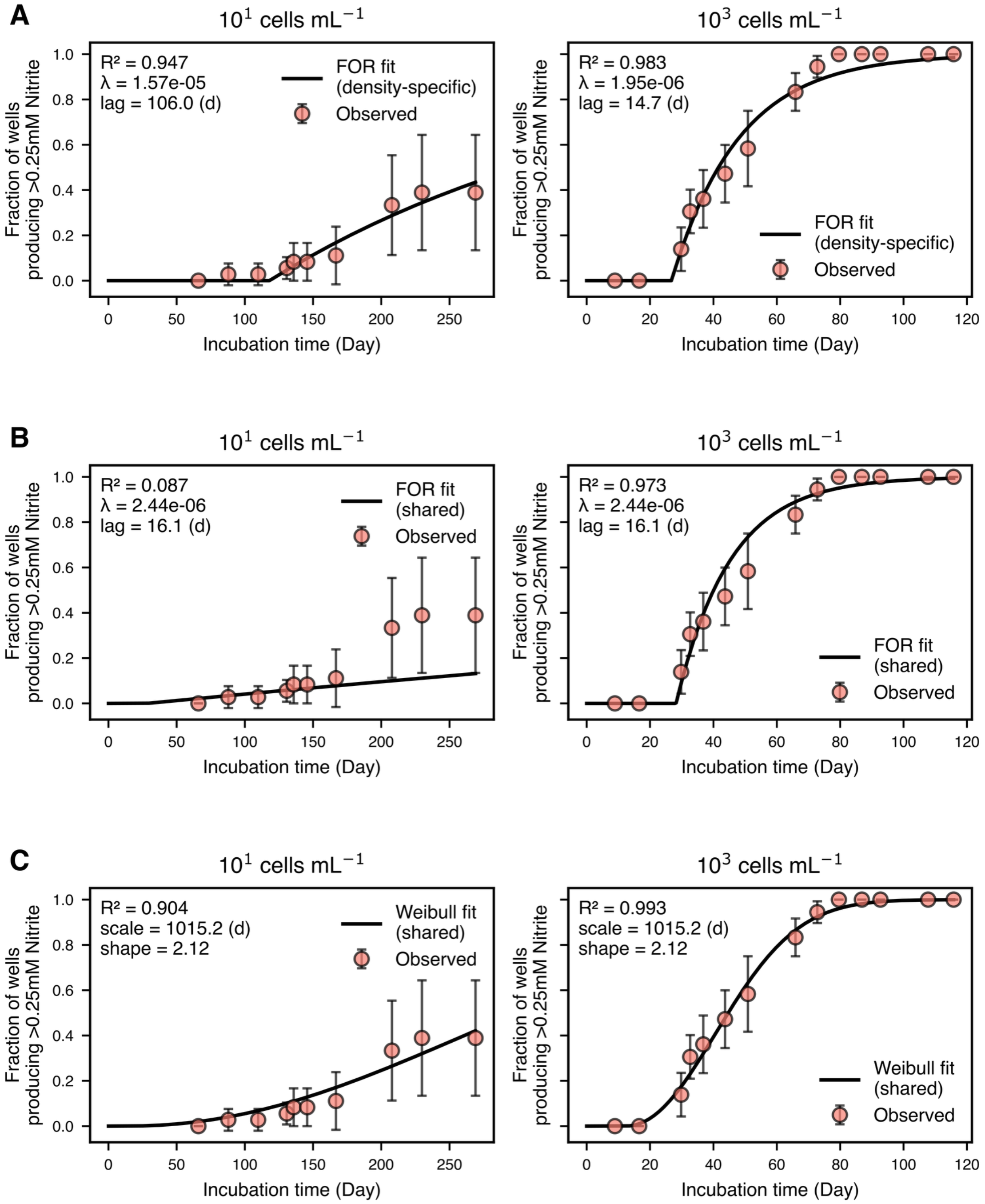


**Figure S14. Fitting of the probability of $\Delta V_t$-independent growth initiation.** The probability of $\Delta V_t$-independent growth initiation was fitted to the time course of the proportion of nitrite-positive wells in batch culture experiments of PY1 in the absence of CFS at low initial cell densities ($10^1$ and $10^3$ cells $mL^{-1}$). Red circles represent experimental data, and black lines represent model fits. The mean values from biological replicates (n = 3) were used as experimental data, and error bars (mean ± SD) are shown. (A) Fitting of the probability of $\Delta V_t$-independent growth initiation using an exponential distribution based on a Poisson process ($F(t) = 1 - e^{-\lambda(t - t_r - lag)}$). Different $\lambda$ values were fitted for each initial cell density. (B) Fitting using an exponential distribution based on a Poisson process with a common $\lambda$ across initial cell densities. (C) Fitting of the shape parameter ($k$) and scale parameter ($\tau$) that govern the probability of $\Delta V_t$-independent growth initiation using a Weibull distribution ($F(t) = 1 - exp\left(-\frac{t-t_r}{\tau}^k\right)$). Common parameters were fitted across initial cell densities; see the Supplementary Methods for details.

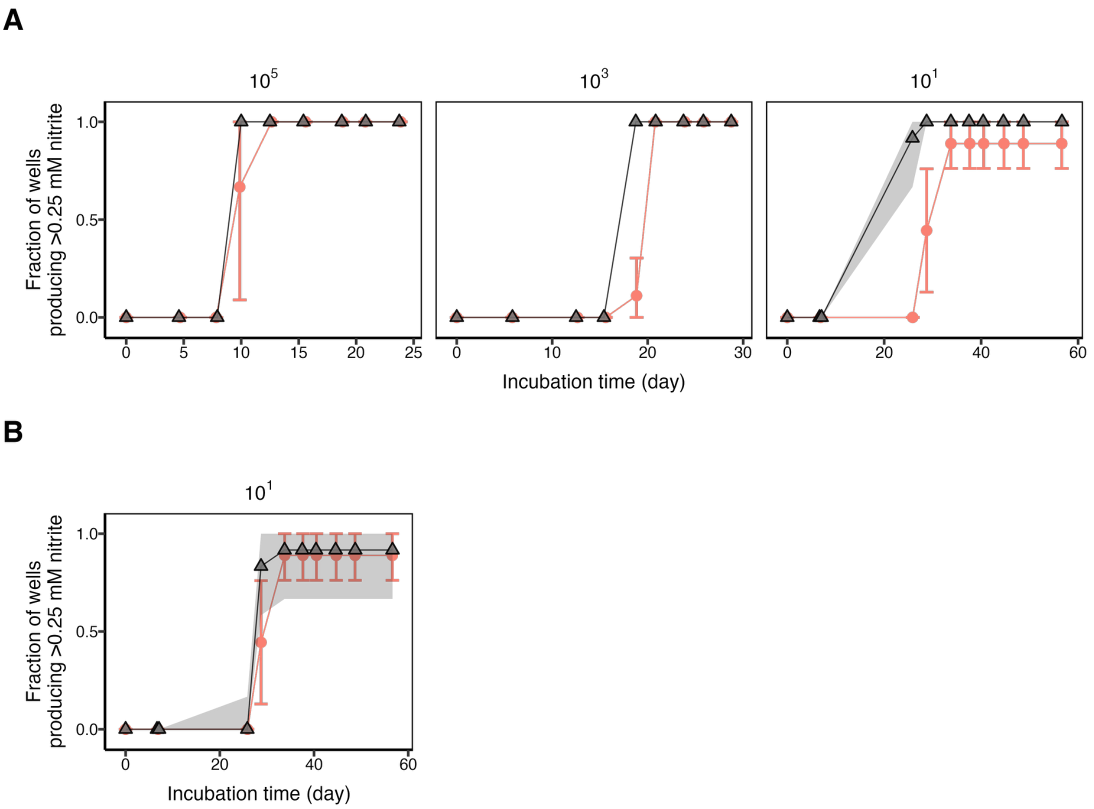


**Figure S15. Simulation of the time course of the proportion of nitrite-positive wells for PY1 in the presence of CFS using a model incorporating $\Delta V_t$-independent growth initiation.** As in Fig. 6C, growth of PY1 in the presence of CFS was simulated using the model incorporating a $\Delta V_t$-independent growth initiation described by a Weibull distribution. Red circles and lines represent experimental data, and black triangles and lines represent simulation results. The initial cell number was stochastically assigned from a Poisson distribution with mean $\lambda$. Simulations were performed 100 times independently, and the median proportion of nitrite-positive wells at each time point is shown. The 95% prediction interval is indicated by a shaded region. (A) Experimental data were overlaid with simulations assuming $\lambda = 10^5$, $10^3$, and $10^1$. (B) Experimental data at an initial cell density of $10^1$ cells $mL^{-1}$ were overlaid with simulations assuming $\lambda = 2.48$.

The time course of the proportion of nitrite-positive wells for PY1 in the presence of CFS was well reproduced using a model incorporating a $\Delta V_t$-independent growth initiation described by a Weibull distribution. As shown in Fig. 6A, a slight discrepancy in the timing of nitrite detection was observed at the initial cell density of $10^1$ cells $mL^{-1}$ (Fig. S15A). This discrepancy may be attributed to the stochastic variability in cell numbers during inoculation and the occurrence of zero-cell wells under low cell density conditions. Therefore, as described in Fig. S12, additional simulations were performed using $\lambda$ estimated from the observed proportion of nitrite-negative wells. When the representative value $\lambda = 2.48$ was used, agreement between simulation and experimental data improved (Fig. S15B). These results indicate that incorporating a $\Delta V_t$-independent growth initiation described by a Weibull distribution does not substantially affect the ability of the model to reproduce the time course of the proportion of nitrite-positive wells in the presence of CFS.

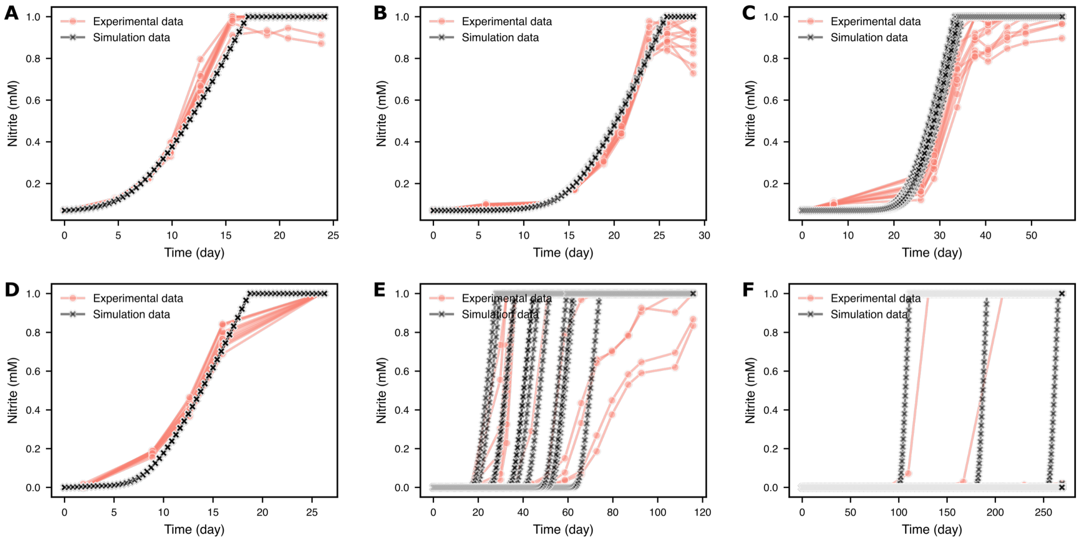


**Figure S16. Time course of nitrite production per well for PY1 in batch culture simulated using a model incorporating $\Delta V_t$-independent growth initiation.** Simulated time courses of nitrite production per well were generated using the same individual-based model (IBM) and parameter settings as in Fig. 6C. Red circles and lines represent experimental data, and black triangles and lines represent simulation results. The initial number of cells was stochastically assigned from a Poisson distribution with mean $\lambda$. Experimental data from a representative biological replicate and the corresponding simulation results are shown. (A–C) Simulation results in the presence of CFS. Experimental data were overlaid with simulations assuming $\lambda = 10^5$, $10^3$, and $10^1$, respectively. (D–F) Simulation results in the absence of CFS. Experimental data were overlaid with simulations assuming $\lambda = 10^5$, $10^3$, and $10^1$, respectively.

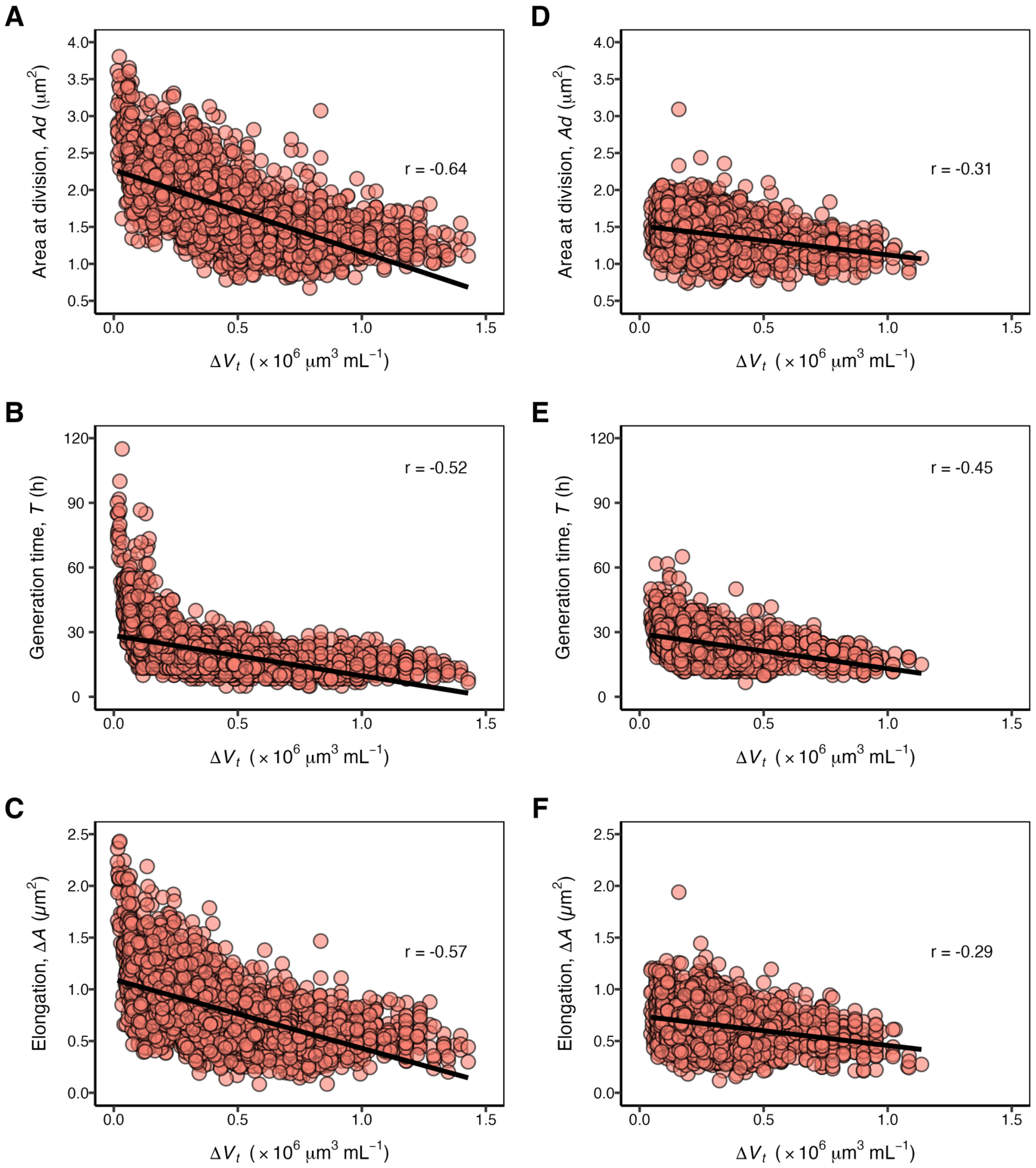


**Figure S17. Comparison of cell size control-related parameters ($Ad$, $T$, $\Delta A$) in PY1 with and without CFS.** Three models of cell size control in bacteria (the Sizer, Timer, and Adder models) suggest that cell division occurs when cells reach a defined size ($Ad$), generation time ($T$), or cell elongation until division ($\Delta A$), respectively [17]. Based on the single-cell observation results of PY1 in this study, the differences in these three parameters between the condition without CFS (A–C) and the condition with CFS (D–F) were visualized. (A, D) Area at division ($Ad$) as a function of $\Delta V_t$ at cell birth. (B, E) Generation time ($T$) as a function of $\Delta V_t$ at cell birth. (C, F) Cell area elongation until division ($\Delta A$) as a function of $\Delta V_t$ at cell birth. A linear regression line fitted to all data is shown to indicate the overall trend. Three FOVs were analyzed for each time-lapse observation experiment, and data from biological replicates (n = 3) were combined.

In the absence of CFS, PY1 showed increases in all three cell size control-related parameters ($Ad$, $T$, and $\Delta A$) under low $\Delta V_t$ conditions. By contrast, in the presence of CFS, the increases in these three parameters under low $\Delta V_t$ conditions were suppressed. This indicates that, in the absence of CFS and under low $\Delta V_t$ conditions, PY1 delayed cell division even after exceeding the cell size and generation time at which division occurred in the presence of CFS. These results suggest that the induction of cell division in PY1 may be suppressed in the absence of CFS.

### Legends for Movies S1–S4

**Movie S1. Representative field of view (FOV) from single-cell observations of PY1 in the absence of CFS.** Time-lapse images from the start of observation (0 h) to the end (163 h 20 min) are shown.

**Movie S2. Representative field of view (FOV) from single-cell observations of *N. europaea* in the absence of CFS.** Time-lapse images from the start of observation (0 h) to the end (32 h 30 min) are shown.

**Movie S3. Representative field of view (FOV) from single-cell observations of PY1 in the presence of CFS.** Time-lapse images from the start of observation (0 h) to the end (101 h 40 min) are shown.

**Movie S4. Comparison of single-cell observations of PY1 in the absence and presence of CFS.** Observations in the absence of CFS were trimmed to match the observation period in the presence of CFS (0 h to 101 h 40 min), and both conditions are displayed side by side.
